# A supervised data-driven spatial filter denoising method for speech artifacts in intracranial electrophysiological recordings

**DOI:** 10.1101/2023.04.05.535577

**Authors:** Victoria Peterson, Matteo Vissani, Shiyu Luo, Qinwan Rabbani, Nathan E. Crone, Alan Bush, R. Mark Richardson

**Author notes:** **Corresponding author:** Correspondence to Victoria Peterson or Alan Bush. Equal contribution.

## Abstract

Neurosurgical procedures that enable direct brain recordings in awake patients offer unique opportunities to explore the neurophysiology of human speech. The scarcity of these opportunities and the altruism of participating patients compel us to apply the highest rigor to signal analysis. Intracranial electroencephalography (iEEG) signals recorded during overt speech can contain a speech artifact that tracks the fundamental frequency (F0) of the participant’s voice, involving the same high-gamma frequencies that are modulated during speech production and perception. To address this artifact, we developed a spatial-filtering approach to identify and remove acoustic-induced contaminations of the recorded signal. We found that traditional reference schemes jeopardized signal quality, whereas our data-driven method denoised the recordings while preserving underlying neural activity.

## Introduction

Human intracranial recordings, i.e., in-vivo electrophysiological signals acquired during specific neurosurgical treatments such as focal epilepsy and deep brain stimulation (DBS), have enabled the study of neural responses with high temporal and spatial resolution, in both surface and deep structures of the human brain during behavioral tasks^1^. The study of speech motor control especially benefits from awake intraoperative recordings during which local field potentials (LFP) and single unit activity of subcortical targets can be simultaneously acquired during speech production^2,3^.

To develop brain-computer interfaces (BCI) for speech protheses, the neural activity in the high gamma band (60-200 Hz) is typically used for speech decoders^4–7^. Recently, we and others have shown that brain activity measured with invasive recordings may contain artifacts associated with the mechanical vibrations produced by the participant’s voice or sounds played by a loudspeaker^8,9^. For overt speech experiments, this speech artifact shares spectral characteristics with the produced audio signal, being locked at the fundamental frequency (F0) of the participant’s voice^8^. The overlap between typical human F0 (between 70 and 240 Hz) and high-gamma activity (60 to 250Hz) imposes the need to account for this speech artifact to study the brain activity associated with speech production.

As shown before, the acoustic artifact is introduced along the acquisition chain, where the mechanical vibrations of the acoustic signal are translated into voltage^9^. Passive electrical components can exhibit an electrical response when stressed physically, a phenomenon referred to as the *microphonic effect*^10^. This effect can be exacerbated in the case of speech tasks performed during stereotactic neurosurgery, at which the patient’s head is fixed to a stereotactic frame (Fig. 1a). This frame may act as a resonance system that exacerbates speech-induced vibrations originating in the larynx and travelling through the head and skull. Speech-induced vibrations, which look like a distorted version of the speech audio, can affect the electrodes and acquisition chain appearing in the neural recordings tracking the fundamental frequency of the participant’s voice (Fig. 1b). Acoustic-induced vibration artifacts can be detected by measuring the coherence value between the speech acoustic signal and neural recordings in the high-gamma frequency band^8^ (Fig. 1c). This coherence value varies largely between and within patients, indicating that the speech artifact is a channel-specific type of noise, and thus traditional re-reference schemes may jeopardize the quality of the neural recordings^11^.

**Fig. 1.**
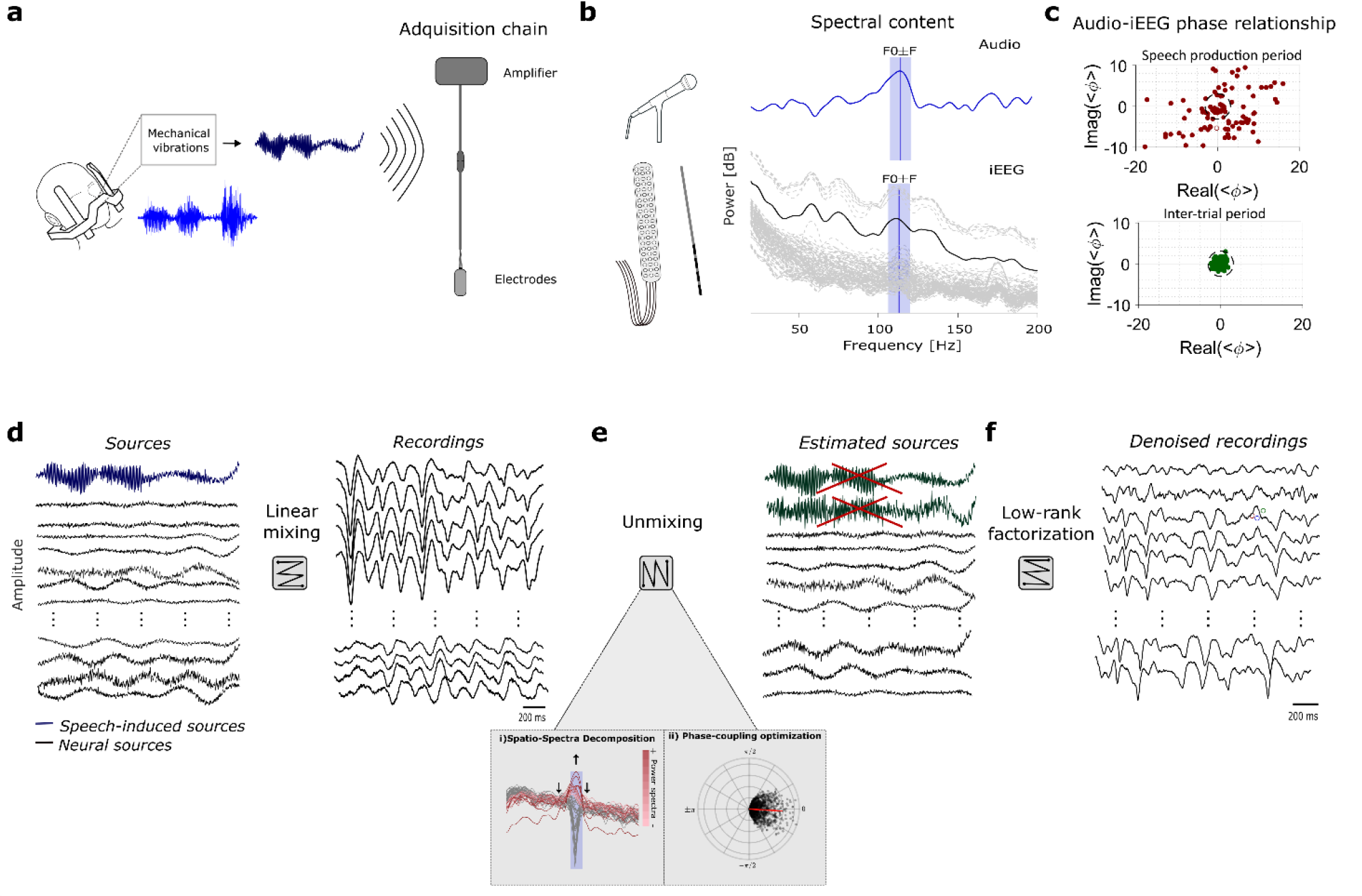
Speech-induced speech artifact model assumptions. **a**, Schematic representation of how speech-induced mechanical vibrations can affect neural recordings during stereotactic frame surgery. The frame attached to the patient’s head acts as a resonance system with the skull, transmitting and distorting speech-induced vibrations that can affect the neural recordings’ acquisition chain. **b**, The speech artifact tracks the fundamental frequency (F0) of the participant’s voice. **c**. The coherence between the audio signal and the neural recordings across trials (< *ϕ* >) can assess the level of speech artifact contamination in each electrode. During speech production many channels present artifact (lie outside the dashed circumference), while no artifact is present during inter-trial periods. **d**, The recordings, at the amplifier level, can be thought as a linear mixing between the speech artifact and brain-related sources. **e**, The proposed unmixing pipeline is based on two spatial-filtering methods. While the Spatio-Spectra Decomposition (SSD) helps to enhance the signal-to-noise ratio in the frequency band of interest, the Phase-coupling Optimization seeks at finding the sources with the highest coherence value with respect to the audio. **f**, By means of low-rank factorization, the signal reconstruction is done zeroing out the speech artifactual sources.

Considering the recorded brain activity as superpositions of different statistical sources^12^, the brain signals at the amplifier level can be thought of as the consequence of a linear mix between true neurophysiological and non-neural sources, including speech-induced vibrations (Fig. 1d). Using spatial filter methods, these sources can be untangled and estimated from the multivariate (multichannel) electrode signals^13^. As such, traditional re-referencing schemes used in neurophysiology, like bipolar, Laplacian, or common average reference (CAR), can be reframed as spatial filtering approaches, in which the recorded brain signals are multiplied by a predefined matrix that references the recording of one electrode with respect to a neighborhood of channels^14^ (see CAR as a spatial filtering algorithm, Methods).

Data-driven spatial filters offer a more flexible re-reference scheme than traditional methods. They can be used for denoising, using linear transformations to estimate the data subspace related to the “noise” and discard it for the subsequent signal analyses^14–16^. This approach has been used primarily for non-invasive electrophysiology^14,15,17,18^, and more recently for iEEG signal processing^13,19,20^. A typical pipeline for artifact removal in non-invasive EEG consists of using principal component analysis (PCA) and independent component analysis (ICA) for identifying artifact components^17,21^. PCA is commonly used as a dimensionality reduction tool to alleviate the decomposition done by ICA. Then, by means of low-rank factorization, backward and forward projections are made between the signal and the source space, identifying and discarding those components related to artifacts. However, due to the nature of the speech-induced vibration artifact, which overlaps in frequency with high-gamma activity, ICA may fail to decompose artifactual from neural sources^22^ (see also Results Section).

Here, we introduce phase-coupling decomposition (PCD), an algorithm for acoustic-induced artifact rejection. This algorithm performs data-driven spatial filtering denoising based on low-rank factorization. It is designed to separate acoustic-induced artifactual sources from neural sources via a phase-coupling decomposition. The spatio-spectral decomposition (SSD)^22^ algorithm is used first to enhance signal-to-noise ratio around F0 and perform dimensionality reduction (Fig. 1e). The phase-coupling optimization (PCO)^23^ method is then applied to identify sources phase-locked to the acoustic signal (Fig. 1e). Thus, the coherence between the audio and the neural data is optimized, allowing retrieval of those sources related to acoustic-induced noise. Similar to the ICA-based artifact pipeline mentioned above, signal reconstruction is based on low-rank factorization, discarding the detected artifactual components (Fig. 1f).

First, we demonstrate how PCD cleans acoustic-induced artifacts from an affected recording (Fig. 2). Then, we test the denoising performance of this algorithm in simulated data, in which the artifact and the neural sources are artificially generated and mixed. The algorithm successfully recovers the artifactual source, in the time, frequency, and phase domains even when dealing with highly non-stationary audio signals (Fig. 3). Importantly, we demonstrate the algorithm’s ability to denoise while preserving neural data and compare its performance with respect to traditional spatial filtering methods like CAR and ICA (Fig. 4). Finally, we test the PCD denoising algorithm in real acoustic contaminated iEEG data, showing a significant reduction of the extent and number of artifact-affected electrodes while preserving the underlying speech-related neurophysiological response (Fig. 5-7).

**Fig. 2.**
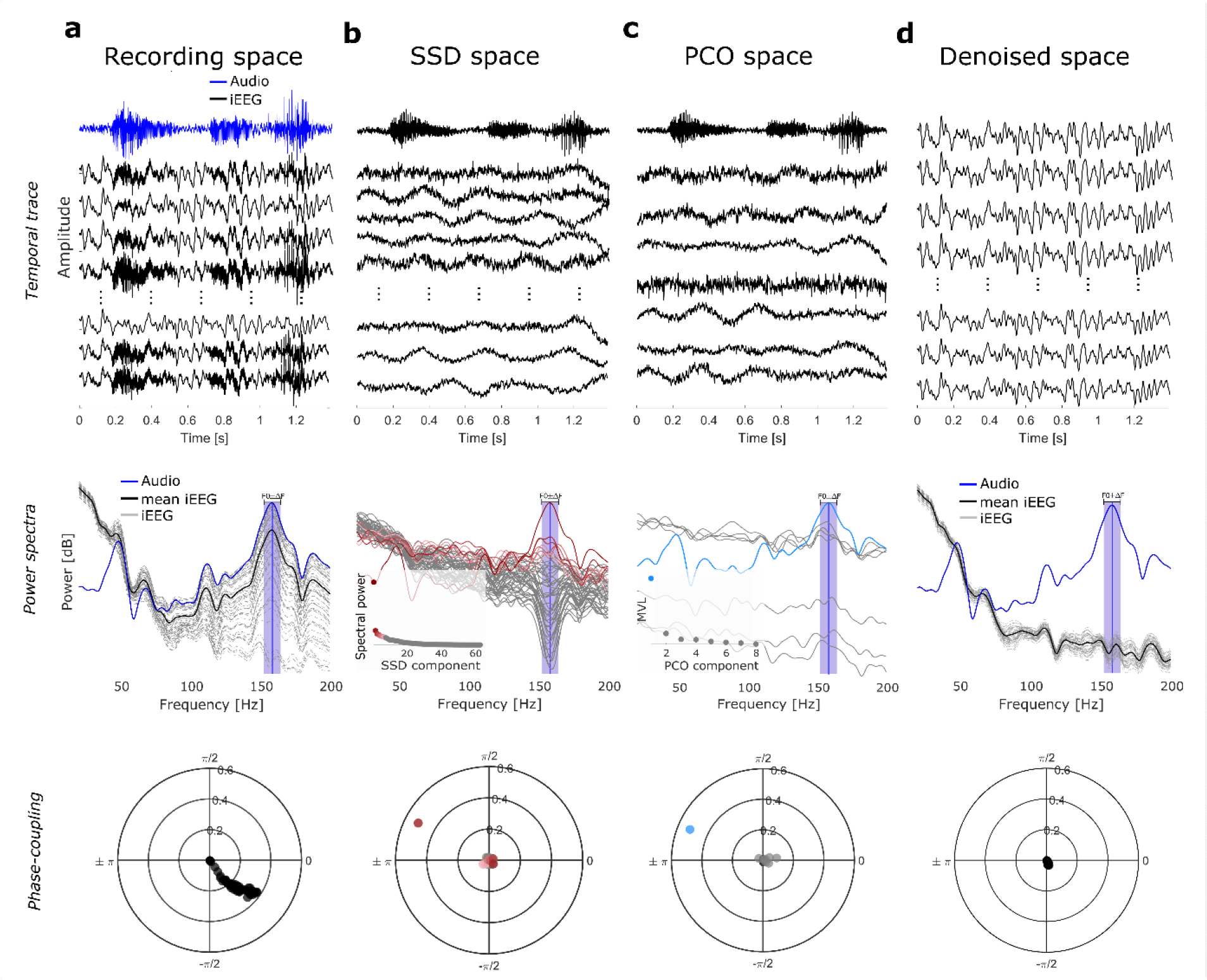
Illustration of the phase-coupling decomposition pipeline. Space transformations from the recording space to the clean space in the temporal, power, and phase domains. **a**, (Recording space), Top: iEEG recordings (black traces) were combined with the recorded audio (blue trace) to simulate the contaminated recordings for illustration purposes. Middle: note the similarity of audio and iEEG power spectra. Bottom: Polar plot showing phase-coupling value between the audio and iEEG channels (represented as dots). **b**, (SSD space), SSD identifies components that maximize power around F0 (color-coded in the spectral domain from dark red to light pink). **c**, (PCO space), The SSD components with the strongest power around F0 are used to compute PCO. The MVL is optimized and only those PCO components that show the highest coherence with the audio are identified as artifactual (light-blue trace). Here, the first 8 components are shown. **d**, (Denoised space), Via low rank-factorization, the artifactual component(s) are excluded for signal reconstruction. Note the clean iEEG traces, with no peak around F0 in the power spectra and phase-coupling values centered at zero.

**Fig. 3.**
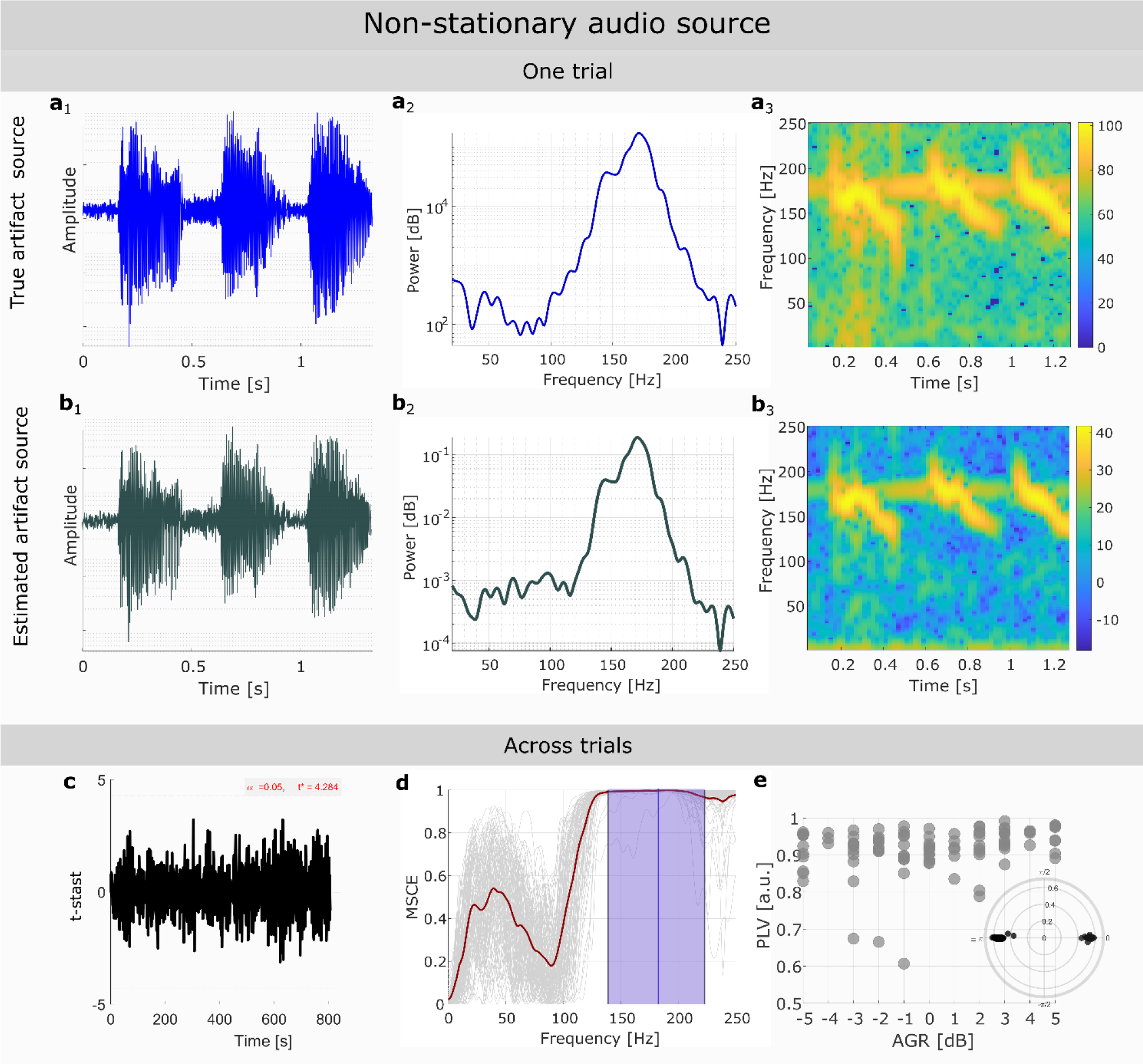
PCD correctly retrieves non-stationary and weak artifactual sources. Representative performance of PCD on an artifactual source with a non-stationary signal profile and an artifact-to-physiological gamma ratio (AGR) = 2dB. **(Top panel) a1-a3**, Visualization in the time, frequency, and time-frequency domain of the ground-truth artifactual source for a given trial. **b1-b3**, Same as above for the retrieved artifactual source for the same trial. **(Bottom panel)**, PCD performance across trials. **c**, One-dimensional statistical parametric mapping was used to evaluate statistical similarities at each sample point between the true and estimated source for each trial. No significant differences (t-stats values < t*) were found at any time point. **b**, Magnitude-squared coherence estimate (MSCE) between true and estimated artifact sources. Gray lines represent individual trials. Mean MSCE across trials is denoted by the dark red line. The mean F0 across trials is shown as the blue vertical line, while the violet band indicates the SAFB. The mean MSCE value in the SAFB was always above 0.97, regardless of the variable AGR across trials. **e**, Phase-locking value (PLV) across simulated trials at different AGR between the true and estimated artifact source, for each trial. Inset on the right corner aggregates the phase difference found across trials between the true and estimated artifact source

**Fig. 4.**
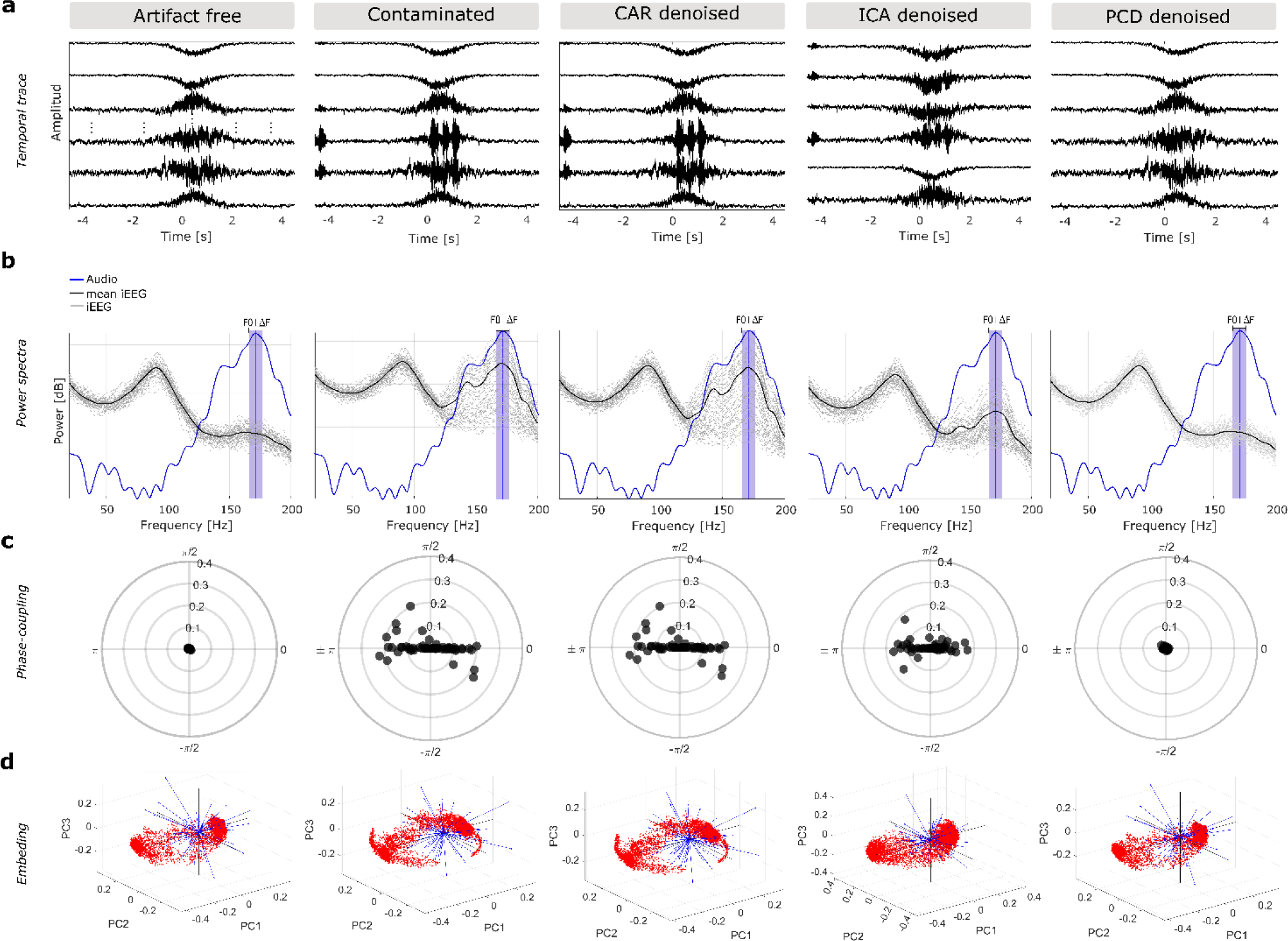
PCD removes acoustic-induced speech artifact while preserving neural activity in simulated data, outperforming previous methods. Impact on signal quality of applying CAR, ICA and PCD for a trial of the realistic simulation (low-amplitude broadband gamma modulation, AGR=-2 dB and non-stationary artifact source). Temporal trace (a), power spectra (b) and phase-coupling plots (c) as in Figure 2. **a**, Resulting temporal traces with time reference relative to the speech onset (t=0). **b**, Power spectrum of each resulting iEEG signal (gray lines) and the recorded audio (blue line). The mean power spectra across iEEG channels is shown by the thick black line. Violet shades indicate the SAFB. **c**, The phase relationship between the audio and the resulting brain signals. Each dot represents a channel. **d**, biplots of the PC subspace described by the first three principal components (PC1, PC2, PC3). Red dots represent scores while blue lines are the loading directions. The shape of the red cloud illustrates the PCA embedding.

**Fig. 5.**
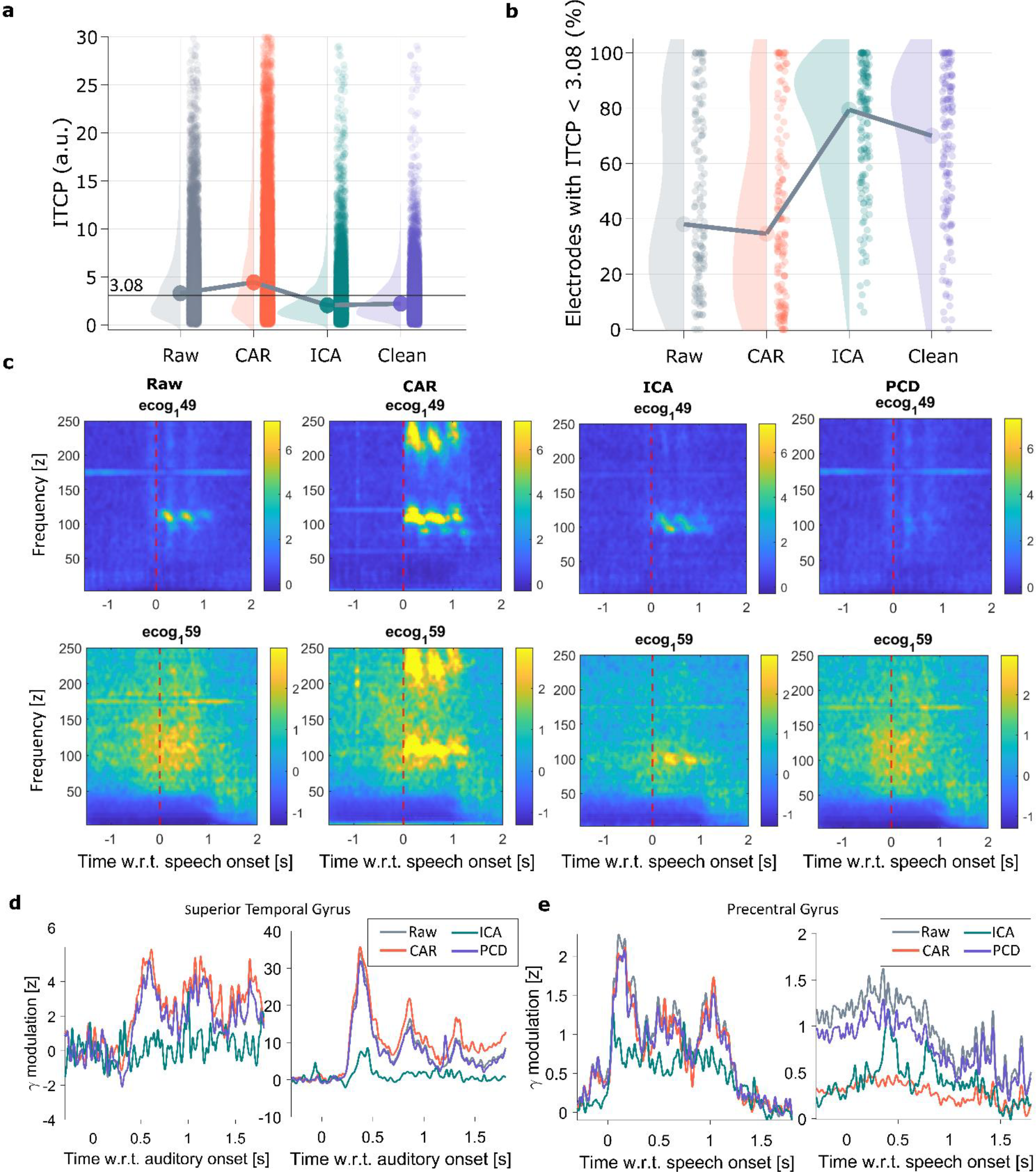
PCD removes the speech artifact while preserving underlying γ-activity in intracranial recordings. **a**, Coherence value between a channel and the recorded audio, measured by means of ITPC. Here a dot represents a channel, and the horizontal solid line indicates the significance ITCP threshold (3.08). **b**, Percentage of electrodes that lie below the significant ITCP threshold, where every dot represents the data from a recording session. **c**, Time-frequency plots for raw signals and effect of applying each denoising method in an electrode with strong speech artifact (top row) and an electrode with physiological speech-related γ-activity (bottom row). Time is relative to speech onset. **d**, Gamma profiles (z-scored normalized) during stimulus onset of two electrodes located in the superior temporal gyrus for two different patient’s data. **e**, Gamma profiles (z-scored normalized) during speech onset of two electrodes located in the precentral gyrus for two different patient’s data. Different color traces are used for the raw and the resulting denoised data via each tested method.

**Fig. 6.**
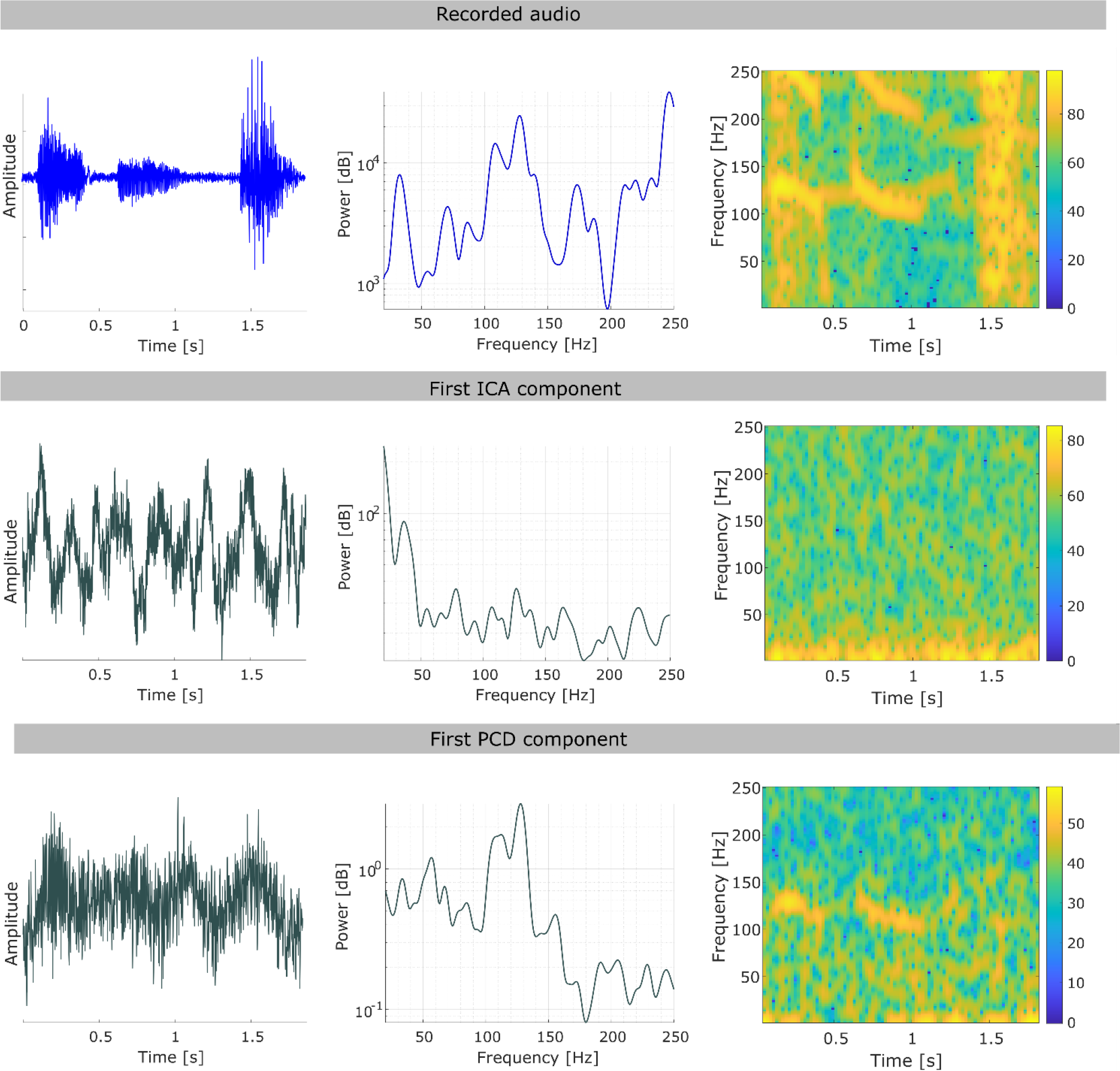
Artifact retrieval comparison between ICA and PCD. **a**, distribution of the number of removed component by each method. **b**, Comparison in the temporal, frequency, and time-frequency domain of the recorded audio and the first artifactual component estimated by ICA and PCD. For a given participant, at a randomly selected trial, the component with the highest phase-relationship with the audio is shown here.

**Fig. 7.**
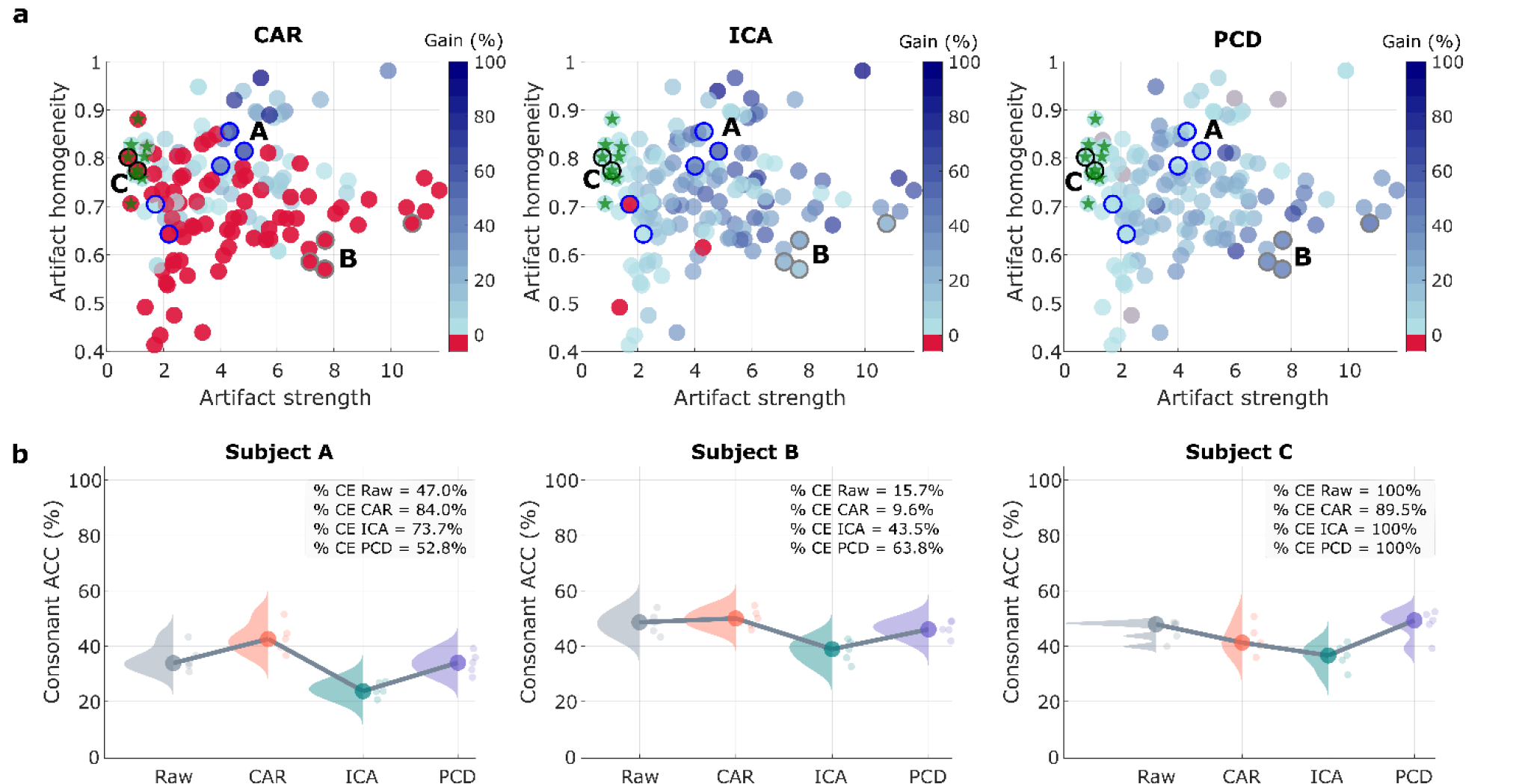
Understanding the impact of each denoising method as a pre-processing step. **a**, Percentage gain in electrodes below the significant ITCP threshold (Gain) versus artifact homogeneity and strength, for CAR, ICA, and PCD. Each dot represents the data from a recording session. Raw data with no detectable speech artifact is marked with a green star. Data from 3 subjects (A, B and C) are highlighted: Homogenous artifact case (Subject A); Strong artifact case (Subject B) and No artifact case (Subject C) represented as circles with blue, gray and black contours, respectively. **b**, Impact in speech decoding from neural data (detection of consonant-vowel syllables) in three cases: i) clean electrode (CE) gain is higher for CAR and ICA than PCD (Subject A), ii) CE gain is better with PCD than for the other two methods (Subject B), and iii) data without speech artifact (Subject C). ACC stands for accuracy, i.e., the number of correct classified samples over the total of samples. Every point represents the accuracy of the testing data on a 5-fold cross-validation scenario.

## Results

### Overview of phase-coupling decomposition

Here, we provide a summary of the PCD pipeline (Box 1, Methods). We consider the brain recordings as a linear combination of statistical sources, where at least one of those sources is related to the speech artifact, which can be detected by measuring the coherence value between the audio signal and the neural recordings^8^. The recorded brain signals then are considered as terms of a linear forward model, in which *N*_*C*_ statistical sources are projected into *N*_*C*_ channels along the *N*_*S*_ data samples via a linear mixing process:

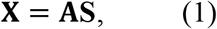

where 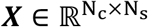 denotes the recorded brain signals, 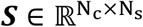 is the source activity, and 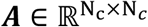 denotes the mixing matrix. Each column ***a*** of the mixing matrix ***A*** is known as *spatial pattern* and describes the projection of an individual source to the sensor space.

The idea of PCD is to find the artifactual sources that are phase-coupled with the acoustic signal, denoted here as 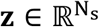. The first step within the PCD pipeline is to estimate the speech artifact frequency band (SAFB). To this end, since the vibration speech artifact tracks F0, the power spectrum of the acoustic signal is used (Step 1 Box 1). Once SAFB is estimated, the spatio-spectral decomposition (SSD) algorithm (Methods)^22^ is applied to (i) enhance the power around the SAFB and (ii) reduce the dimensionality of the data (Step 2 Box 1).

#### Box 1

**Phase Coupling Decomposition**

Given the recorded brain activity 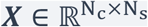, where *N*_*c*_ and *N*_*s*_ are the number of channels and samples, respectively, and an acoustic signal 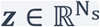, this algorithm identifies narrow-band components of ***X*** that are phase-coupled with ***z***.

**Step 1:** Estimate the speech artifact bandwidth.

1. Calculate the power spectrum of ***z***.
2. Around the fundamental frequency (F0) of the voice, find the peak in the spectra (*P*_*F0*_).
3. Define the center *P*_*F0*_ and bandwidth of the speech artifact frequency band (SAFB): *F*_*c*_ ± Δ *F*_*c*_.

**Step 2:** Find projections of ***X*** that maximize the signal-to-noise-ratio (SNR) around *F*_*c*_ via the spatio-spectral decomposition^22^ (SSD).

1. Define as *signal band* the SAFB: *F*_*c*_ ± Δ *F*_*c*_, and as *noise band all frequencies* except SAFB. Find the projection matrix ***W***_*SSD*_ by solving equation (3) and project the data on the first *k* SSD components 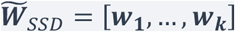, as follows: 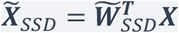.

**Step 3**: Find the artifactual sources via the phase-coupling optimization algorithm^23^ (PCO).

1. In the SSD space, find the projection matrix ***W***_*PC0*_ which maximizes the mean vector length (see equation (6) and (7)) between 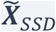 and ***z***.
2. Compute the PCO components 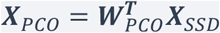 and identify the *m* acoustic-induced sources.

**Step 4:** Perform the denoising via low-rank factorization.

1. Compute the PCD unmixing matrix ***W***_*PCD*_ which projects ***X*** to the artifact source space. This matrix is the conjunction of both ***W***_*PC0*_ and ***W***_*SSD*_ (equation (8)).
2. Compute the PCD mixing matrix via the pseudoinverse:

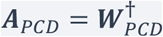
3. Reconstruct the signal by using (*N*_*c*_ − *m*) components: 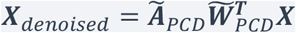.

See details in the Methods.

Data projected onto the first *k* SSD components, showing the largest power at the SAFB, are used for the phase-coupling optimization (PCO) algorithm^23^ (Methods), in which an index of coherence between the data on the SSD space ***X***_*SSD*_ and the acoustic signal ***z*** is maximized (Step 3 Box 1). Following previous work^20^, we refer to this index of coherence as the Mean Vector Length (MVL, see Methods equation (6)). Note that phase-coupling is calculated across sample points, such that MVL represents a summary of the phase relation across a given time window. The components showing highest coherence with the acoustic signal are considered as speech artifact sources and will be excluded during signal reconstruction (Step 4 Box 1).

Fig. 2 depicts how contaminated recordings are denoised by applying PCD, in the temporal, spectral and phase domains. For illustration purposes, we artificially contaminated iEEG recordings with an audio signal (Recording space, Fig. 2a), resulting in iEEG signals with a strong peak in spectral power around F0 (Fig 2a center), together with a consistent phase-relationship with the acoustic signal (Fig 2a bottom). After SSD is applied, only a few components have enhanced power spectra around F0 (Fig. 2b center). Applying PCO in the SSD space gives us components maximally phase-coupled with the recorded audio (Fig. 2c bottom). The components identified as artifactual will be discarded for signal reconstruction, thus achieving a denoised signal (Fig. 2d).

### Method benchmarking on in-silico data

To benchmark our method’s denoising performance we applied PCD on simulated neural data with added simulated speech artifact. This approach allows direct comparison of denoised data to the ground-truth simulated neural signals. Briefly, we simulated recurrent networks of leaky integrate-and-fire (LIF) neurons (N = 5000, 80% excitatory and 20% inhibitory) to simulate 100 physiological broadband γ-source activities, defined as the summation of absolute values of the excitatory and inhibitory currents entering the excitatory neurons population (Supplementary Fig. 1, Methods). The model captures key features of physiological LFPs, including(i)the entrainment of the low-frequency fluctuations to the external input, (ii) the internal resonance of broadband γ-oscillations driven by the mean rate of the external input (Supplementary Fig. 2) and(ii) phase-amplitude coupling between low and high frequency oscillations (Supplementary Fig. 3). We defined a single simulated speech artifact source assuming that it is identical to the produced audio signal, denoted as ***z*** in the PCD pipeline. After adjusting the artifact-to-physiological gamma ratio (AGR), we linearly projected the sources to the simulated recordings by the application of a random mixing matrix. We tested the PCD pipeline in different simulation scenarios based on the expression of the external input driving LIF neurons and the audio source (Supplementary Fig. 1, Methods). Toy scenarios were created to stress the method under different pre-defined source conditions (Methods, Supplementary Fig 4-8.)

To test the method under realistic conditions, we modulated the external input to the neural source to mimic observed γ-band activity during 60 trials of a speech production task (Supplementary Fig. 9). Neural sources were mixed (at a range of AGRs) with recorded audio of Parkinson’s Disease patients during an intraoperative speech task (Supplementary Fig. 10). PCD reliably recovered the artifactual source, as can be observed in the time, frequency, and time-frequency profiles of both the true (Fig. 3a) and estimated (Fig. 3b) artifact source. Across trials, no differences were found at any sample point between the true and the estimated artifact source, as assessed by one-dimensional statistical parametric mapping (SPM1d, https://spm1d.org/), (Fig. 3c). The estimated artifactual source had high coherence with the true artifactual source at frequencies around F0 (mean MSCE > 0.97, Fig. 3d), independently of differences in the AGR (Supplementary Fig. 11). Finally, the estimated source was either in phase or anti-phase relationship with respect to the true source (Fig. 3e). Similar results were found for the other realistic simulations (Supplementary Fig. 11).

### PCD removes the speech artifact while preserving simulated γ-activity

Next, we compared the performance of PCD with CAR and ICA, two well-known preprocessing spatial filtering approaches, using simulated data under the realistic scenario (see Methods). For a fair comparison, all three methods were implemented on a trial-wise basis and components identified by ICA were scored according to their phase-coupling value against the recorded audio (Methods). We evaluated performance in terms of the capacity of each method to retrieve the temporal, frequency, and phase information of the simulated ground-truth neural data (Fig. 4a-4c). Figure 4 shows the simulated neural activity without the artifact source (Artifact free), when linearly combined with the artifactual source (Contaminated), and the resulting denoised signals from each method (CAR, ICA, PCD). PCD outperformed the other methods in terms of its ability to clean the data and retrieve the simulated brain signals, as can be observed in the time (Fig. 4a), frequency (Fig. 4b) and phase (Fig. 4c) domains. In this simulation CAR produced traces very similar to the contaminated signal, while ICA attenuated the artifact to some extent. Interestingly, PCD completely removed the narrow-band component induced by the speech artifact while preserving gamma modulation observed in the simulated brain signals.

To assess the preservation of physiological brain signals after denoising, we compared the neural embedding defined in the subspace spanned by the first three PCA components (PC subspace). Note how the PC embedding is distorted in the contaminated signals, as compared to the artifact-free signals (Fig. 4d). CAR resulted in an embedding indistinguishable from the contaminated signals (Fig. 4d), consistent with the time, frequency, and phase analyses. Interestingly, while ICA was able to attenuate the speech artifact, the PC embedding was different from that for the artifact-free signals, indicating that ICA distorted the underlying physiological sources. Here, PCD was the only algorithm that completely removed the artifactual source, as assessed by the signals’ power spectrum (Fig. 4b) and phase coupling (Fig. 4c) plots, while simultaneously preserving the underlying physiological activity, as evidenced by the indistinguishable PC embedding to that of the artifact-free signals (Fig. 4d). To quantitively assess the similarities to the ground-truth PC embedding we used the cosine similarity metric of the three first components (see Methods equation (12)). The mean resulting value was of 0.79, 0.74 and 0.99 for CAR, ICA and PCD, respectively, while for the contaminated data this value was 0.79.

### Denoising PCD performance in acoustic contaminated iEEG data

We applied the PCD pipeline to intracranial recordings of 54 patients undergoing DBS implantation surgery who participated in an intraoperative speech task (Methods). Participants were instructed to repeat consonant-vowel syllables triplets played on earphones. Each participant performed between 1 to 4 recording sessions of up to 120 syllable triplets each. High-density electrocorticography (ECoG) from left ventral sensorimotor cortex, superior temporal gyrus and inferior frontal regions were acquired simultaneously with subcortical recordings. Microelectrode recordings (MER) and LFPs from macro rings located 3mm above each microelectrode tip were registered from the subthalamic nucleus (STN) or globus pallidus internus (Gpi). LFPs were also registered from DBS lead located either in the STN, Gpi or the ventral intermediate nucleus of the thalamus (VIM). As we previously described, around 40% of channels in this dataset show speech-induced vibration artifact^8^.

Considering that the source of the speech artifact is the same across different types of electrodes, the denoising pipeline was applied to all available iEEG recordings together (LFP from ECoG, the DBS lead, and the ring and tip contact of the microelectrode). Given that phase relationships between the recorded audio and the neural recordings are not consistent across trials (Supplementary Fig. 12), data cleaning occurred in a trial-wise manner. The resulting denoised signals were compared to those obtained after applying CAR and ICA as in Fig. 4 (Methods). The number of components to be removed was automatically selected based on the elbow-detection on the MVL values.

To assess denoising performance we used the distribution of Inter-Trial Phase-Consistency with the audio (ITPC, see Methods equation (13)), (Fig. 5a) and the percentage of electrodes without significant ITPC (Fig. 5b) before vs. after applying a denoising framework (Raw vs. CAR, ICA, PCD). Results show that CAR exacerbated the artifact, shifting the ITPC distribution towards higher values (Fig. 5a), whereas ICA and PCD reduced ITPC values and increased the percentage of electrodes without significant coherence with the audio (Fig. 5a-b). However, as shown next, the low coherence values for ICA were due to an aggressive removal of all high frequency components (artifactual and physiological).

To illustrate the method’s performances, time-frequency plots are shown from two electrodes of the same participant and trial. Electrode ecog_1_49 has a strong artifactual narrow-band component after speech onset (Fig. 5c top panel), while ecog_1_59 has a characteristic physiological gamma modulation around the time of speech onset (Fig. 5c bottom panel). Note that the speech artifact is either exacerbated or artificially introduced after applying CAR due to the presence of the artifact on other electrodes (Fig. 5c – CAR). Interestingly, ICA abolished the physiological gamma modulation observed in ecog_1_59, whereas PCD preserved this activity while simultaneously removing the narrowband component (Fig. 5c).

It is of particular interest to analyze the high-gamma profiles before and after applying each denoising method. To this end, we extracted high gamma power time-locked to the auditory stimulus onset and the speech onset, for electrodes located in auditory cortex (superior temporal gyrus, Fig. 5d) and motor cortex (precentral gyrus, Fig. 5e), respectively. The gamma response shown in the auditory cortex during auditory stimulus presentation was unaffected by the artifact, however it was greatly attenuated by ICA (Fig. 5d). Similarly, ICA also abolished the gamma response during speech production for the electrodes shown over motor cortex (Fig. 5e).

To further explore the differences in performance across ICA and PCD, we studied the artifactual sources retrieved by these two methods, as well as the number of removed components. Although on most of the cases between 1 and 4 components were removed by both ICA and PCD (Supplementary Fig. 13), the artifactual sources retrieved by ICA does not resemble a speech-induced artifact (Fig.6).

Additionality, to explore if the variability in denoising performance can be explained by characteristics of the artifact across electrodes, we evaluated the relationship between the relative gain of electrodes (i.e., reduction of percent of electrodes with significant coherence with the audio) by each denoising method, artifact strength and artifact homogeneity (an ITPC-based measure of how similar the artifact is across electrodes, see Methods equations (14) and (15)). Results are shown in Fig. 7a. Note that recordings with negative gain (i.e. increased in percentage affected electrodes) are shown in red. CAR has good denoising performance only for highly homogeneous artifacts with a mild to moderate artifact strength (Pearson correlation ***r***_*gain*−*homogeneity*_ = 0.59, p < 0.0001; ***r***_*gain*− *strength*_ = 0.23, p = 0.006) and can outperform the other methods under these conditions (e.g. Subject A, blue line circles in Fig 7a). Note that CAR can introduce artifact to clean data (e.g., subject C, black line circles in Fig 7a). For ICA, stronger artifacts with mild to moderate homogeneity resulted in higher gain of clean electrodes, as assessed by ITPC (***r***_*gain*−*strength*_ = 0.57, p < 0.0001; ***r***_*gain*−*homogeneity*_ = 0.24, p = 0.003). Interestingly, PCD gain also increases with higher artifact strength but is not significantly correlated with artifact homogeneity (***r***_*gain*−*strength*_ = 0.50, p < 0.0001; ***r***_*gain*−*homogeneity*_ = −0.12, p = 0.14), as illustrated by Subject B (gray line circles in Fig. 7a).

Finally, we assessed the effect of applying each denoising method as a pre-processing step on decoding performance of a densely-connected convolutional neural network (DenseNet)^24^ for consonant decoding (Methods). We tested the decoding method in three different cases: i) CAR and ICA gain is greater than PCD gain (Subject A), ii) PCD outperforms CAR and ICA (Subject B), and iii) data has no artifact (Subject C). Fig. 7b shows that consonant classification accuracy is similar or better when PCD is applied as compared to classification on raw data. Conversely, ICA always decreases classification accuracy, despite increasing the number of electrodes bellow the significant ITPC threshold. Interestingly, for Subject A although the number of contaminated electrodes increased after CAR, the consonant classification accuracy improved, suggesting that decoding capacity in this subject might be partially driven by the artifact (Fig. 7b -Subject B).

## Discussion

Speech-induced vibrations affect the quality of iEEG signals recorded during speech production. This recently described acoustic-induced artifact^8,9^ overlaps in time and frequency with gamma-band activity, potentially compromising the reliability of speech-related analyses, including those used in BCI development. In this paper, we demonstrated that traditional spatial filtering approaches are not appropriate for denoising the speech artifact (Fig. 4-5). Specifically, we showed that CAR exacerbates the presence of the speech artifact when it is heterogeneous across recording channels, “subtracting in” the artifact to otherwise unaffected channels (Fig. 5 - CAR). Although ICA reduces the coherence of neural signals with the audio (Fig. 5a - ICA), this comes at the cost of a strong degradation of physiological γ-band modulations (Fig. 5d,e - ICA), which ultimately results in a reduction of speech-decoding performance from neural data (Fig. 7b - ICA).

In recent years, data-driven spatial filtering methods have been introduced as re-referencing schemes^13,15,25^. Such is the case of SSD, an effective method to increase the SNR for narrow-band components, in which not only the central frequency of the band of interest is enhanced but also its harmonics^13^; a characteristic that is particularly suitable for denoising speech-induced artifacts. Given that the speech contaminations can be assessed by means of coherence with audio channels (ITPC)^8^, PCO (another data-driven spatial-filtering method) is ideally suited to decompose the acoustic artifact source from brain recordings. By combining these methods, we developed PCD, a novel data-driven algorithm for denoising brain recordings contaminated with acoustic-induced artifacts (Fig. 1, Fig.2). Here, we show that PCD can retrieve the speech artifact while preserving physiological γ-band modulations at overlapping frequencies (Fig. 3-4). Through extensive simulations we show that PCD works for different number of channels (from 3 to 100), different ratios of artifact to neural sources amplitude (from -100 to 30 dB), and across different durations of simulated artifact (from 0.5 to 3.5 s), although it is sensitive to the SSD filter parameters like the bandwidth around F0 (Δ*F*) and the filter order (Supplementary Fig. 5,7). Experiments in real data showed that PCD can diminish the number of artifactual electrodes without distorting the underlying neural response (Fig. 5, 7). High-gamma profiles show that PCD (but not ICA) preserves physiological gamma responses (Fig. 5 d,e), indicating PCD represents a more reliable denoising method for these speech artifacts than other traditional pipelines. This finding was also replicated when inspecting the detected artifactual source (Fig. 6) and when testing each denoising method as a pre-processing step of a deep learning network for consonant detection (Fig. 7b).

The PCD method has several underlying assumptions. First, it assumes the speech artifact is a narrow-band component around the fundamental frequency of the participant’s voice, which must be estimated for each participant from audio recordings. Second, it assumes the artifact source is common across electrode modalities, thus allowing the combination of all recording modalities which maximizes the chance of extracting the artifact source^26^. While this might be counterintuitive, the artifact is likely due to mechanical vibrations of cables and connectors along the recording chain^9^ which can affect different recording modalities in the same way. Third, we assume that the recorded audio is a good estimation of the true artifactual source (i.e., there is no spectral distortion or delay between audio and artifact). While this is a strong assumption, currently there is no better proxy for the artifactual source than the recorded audio. Violations of this assumption may explain the difference between simulated (Fig. 3-4) and real data performance (Fig. 5).

The current implementation of PCD has several limitations. (i) Performance declines for broadband artifactual sources (Supplementary Fig. 7), a limitation inherited from SSD, which can enhance the power only in narrow frequency bands^13,22^. This limitation could also partially explain the differences in performance between simulated and real data. (ii) The method does not account for systematic distortions between the recorded audio and the speech artifact. Modelling such distortions might be a promising approach to further improve the method’s performance in future studies. (iii) PCD is computationally expensive given that it involves solving a non-convex optimization problem, thus requiring several runs until it converges to the best solution^23^. As such, PCD for online BCI applications would require further ad-hoc implementations to speed up the optimization process. (iv) The method must be applied in a trial-wise manner given that the artifact’s phase-relationship to the audio strongly varies across trials (Supplementary Fig. 12). For this reason, the method was fitted to individual speech production epochs to optimally estimate the artifact and then a wider window was applied to avoid introducing discontinuities that could affect subsequent analyses. Applying PCD per trial has the additional advantage of reducing memory requirements and computational cost.

Despite the orders of magnitude greater resolution of intracranial recordings compared to scalp EEG signals, the potential for induced speech artifacts is still a significant concern^8,9^. Critical to BCI development using intracranial recordings from overt speech production is the recognition that these artifacts can hamper training of robust machine learning models, which may be biased and fail when used for speech-impaired patients in real-life applications. Signal preprocessing should verify not only that artifactual components are removed or attenuated, but also that neural signals are retained. The PCD method may facilitate the building of BCI decoding models upon reliable neural data by avoiding artifactually driven performance. Additionally, PCD was designed specifically to mitigate loss of signal quality we observed using standard pre-processing methodologies for speech artifact denoising. Using simulated data, we showed that when PCD’s assumptions are satisfied, the method can completely remove the speech artifact while preserving speech-related γ-band modulations. For real data, for which the underlying assumptions might not be strictly satisfied, PCD still achieves significant reductions of the strength and extent of the speech artifact. The consonant decoder results obtained with PCD-denoised signals illustrate the practicability of this method for potential improvements in the reliability of BCI frameworks. Moreover, although this study focused on denoising speech artifacts in intracranial recordings collected intraoperatively, PCD can be used to remove any narrow-band artifact from electrophysiological data, for which an estimate of the source is available.

## List of abbreviations

AGR: Artifact to physiological gamma ratio
BCI: Brain-computer interfaces
CAR: Common average reference
CAS: Colored noise audio scenario
CS: Cosine similarity
DBS: Deep brain stimulation
ECoG: Electrocorticography
EEG: Electroencephalography
F0: Fundamental frequency of the voice
FWHM: Full width at half maximum
ICA: Independent component analysis
iEEG: Intracranial electroencephalography
LFP: Local field potentials
LIF: Leaky integrate and fire
MCAS: Modulated colored noise audio scenario
MSCE: Magnitude squared coherence estimate
MVL: Mean vector length
PC: Principal component
PCA: Principal component analysis
PCD: Phase-coupling decomposition
PCO: Phase-coupling optimization
PLP: Perceptual linear prediction
PR: Participation ratio
RAS: Recorded audio scenario
SAS: Sinusoidal audio scenario
SNR: Signal-to-noise ratio
SSD: Spatio-spectral decomposition
SAFB: Speech artifact frequency band

## Methods

### The phase coupling decomposition algorithm

In the following, we denote the multivariate iEEG signals (acquired using ECoG or LFP either from mapping electrodes or DBS leads), by the matrix 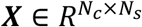, where *N*_*c*_ an *N*_*s*_ represent the number of channels and sample points, respectively. The acoustic signal associated to the speech artifact is denoted by vector 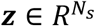. The phase coupling decomposition algorithm (PCD) seeks to find those artifactual sources by solving the inverse problem related to (1) so that each source can be estimated by:

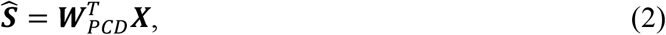

where T denotes transpose (see Box 1). It can be thought of as a supervised two-step spatial filtering procedure which denoises based on low-rank factorization^16^. As shown in Box 1, the speech artifact rejection comprises four main steps: 1) band-width estimation, 2) spatio-spectral decomposition, 3) phase-coupling optimization and 4) signal reconstruction. In the following, each step will be described.

#### 1) Speech artifact band-width estimation

The speech artifact is a narrow-band component noise. It happens around the fundamental frequency (F0) of the participant voice^8^. The recorded audio signal ***z***, is used as a proxy for the speech artifact frequency band estimation. We use Welch’s method^27^ for calculating the power spectra of the audio (detrended signal). An estimate of F0 from audio recordings is used as a starting point to find a peak in gamma band (50-250 Hz). Here, the Gaussian curve fitting algorithm is used to calculate the central frequency (*F*_*C*_) and band width (Δ*F*_*c*_) of the speech artifact frequency band (SAFB) by estimating the mean and full width at half maximum of the resulted fitted signal. The SAFB is then defined as *F*_*c*_ ± Δ *F*_*c*_.

#### 2) Spatio-spectral decomposition

The spatio-spectral decomposition algorithm (SSD) is a spatial filtering approach that allows to maximize the signal power at a given frequency peak while minimizing it for surrounding frequencies^22^. The method assumes that the recorded brain activity is a superposition of narrow band oscillations at the given frequency peak of interest (*signal*) and neighboring frequency bands (*noise*), that is ***X*** = ***X***_*signal*_ + ***X***_*noise*_. By traditional temporal filters, such as the Butterworth zero-phase filter, the signal and noise contributions to ***X*** can be separated^22^.

The objective of SSD is to find a set of spatial filters (columns of an unmixing matrix **W**_*SSD*_) such that the power of the signal is maximized while the power of the noise is minimized:

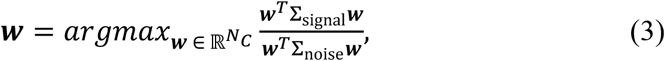

where 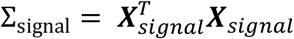 and 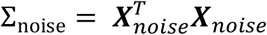are the covariance matrices on the *signal band* and *noise band*, respectively.

The number of components returned by SSD is equal to the number of channels given in ***X***, sorted according to the relative power spectrum in the frequency band of interest. Using the first *k* SSD components, dimensionality reduction can be achieved by projecting the data on the reduced SSD space as follows:

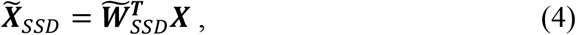

where 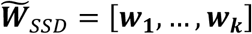.

In the context of the vibration artifact denoising we consider here that our frequency band of interest is the SAFB. Also, the data projected on the first *k* SSD components 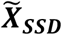 is used as the starting point to run the second spatial filtering technique involved in the PCD pipeline.

#### 3) Phase-coupling optimization

Phase-coupling optimization (PCO) is a supervised spatial filtering algorithm developed by Waterstraat et al. that seeks to recover neural sources that are phase-coupled to an external target variable^23^. The optimization criterion is based on the maximization of the mean vector length^28^ (MVL), a differentiable metric that results in values different from zero when a phase coupling exists. Within this denoising pipeline, we take advantage of the PCO formulation and extend it in order to find the artifactual sources underlying neural activity. Phase coupling is then computed via the MVL between the SSD data projection 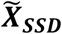 and the best guess of the artifact source. Considering that data in SSD space belong to the complex domain, the analytic signal ***Y***_*SSD*_ ∈ ℂ should be first obtained. The Hilbert transform is utilized to find the imaginary part such that:

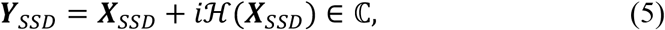

where *i* denotes the imaginary unit defined so as *i*^2^ = −1.

Therefore, the MVL is defined by:

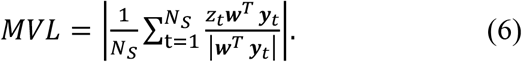

Assuming that the artifact source is the recorded audio signal ***z***, a linear filter ***w***_*PC0*_ = ***w*** = [*w*_1_, …, *w*_*k*_] is found by maximizing the mean vector length at each sample point between the acoustic signal and the SSD components, as follows:

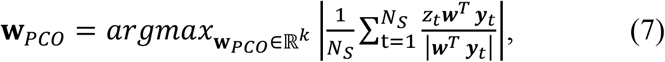

where |·| denotes the amplitude of a complex signal and ***y***_*t*_ is *k* − dimensional vector at the *t*^*th*^ − sample point.

Although the function defined in (7) is differentiable, convexity with respect to ***w*** cannot be guaranteed. Thus, the best solution is selected out of a pool of different solutions found by several random initializations of ***w***. Typically, between 10 and 15 independent runs of the algorithm are done, and the solution with the largest MVL is selected as the best one. The complete set of spatial filters that defines the PCO demixing matrix ***W***_*PC0*_ ∈ ℝ^*k*×*k*^ is found by solving (7) iteratively, outprojecting the previously found filters from the solution space^23,29^, in which data was firstly whitened by a matrix ***M***. The column vectors of ***W***_*PC0*_ are sorted in decreasing order according to the MVL value found during the optimization procedure. Thus, the first *m* components from the resulting data projection in the PCO space 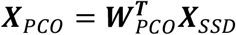 correspond to the speech artifact sources.

#### 4) Signal reconstruction

The objective of the PCD pipeline is to denoise signals contaminated with the speech artifact. The artifact source estimation is just a proxy to facilitate data cleaning. Given that two spatial filtering transformations are applied as a chained process, the PCD unmixing matrix ***W***_*PCD*_ which project the raw data from the amplifier space to the artifact source space should be computed by taking into account the learned linear transformations ***W***_*PC0*_ and ***W***_*SSD*_ as follows:

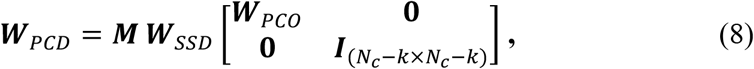

where ***I*** denotes the identity matrix of dimension *N*_*c*_ − *k* × *N*_*c*_ − *k*, and ***M*** is the *N*_*c*_ × *N*_*c*_ whitening matrix applied to find the set of solution vectors to (7).

Once the unmixing matrix is computed, the mixing matrix that defines the forward model explained in (1) can be calculated based on its pseudoinverse:

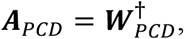

where † denotes the Moore–Penrose pseudoinverse. Then the set of equations that define the linear forward and backward model is given by:

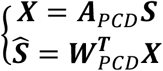

Now, by low-rank factorization the denoised signal can be reconstructed by zeroing from the mixing and unmixing matrix the first *m* components which were previously identified as artifactual. Thus, signal reconstruction is simply done by:

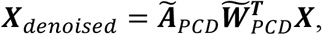

where 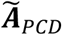 and 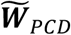 represents the mixing and unmixing matrices of rank *N*_*c*_ − *m*, respectively.

### Simulated Neural Data

#### Description of the neural source model

We simulated a recurrent network of leaky integrate-and-fire (LIF) neurons (N = 5000): 80% excitatory neurons I with AMPA-like synapses and 20% inhibitory neurons (I) with GABA-like synapses^30,31^ (Supplementary Fig 1, Supplementary Table 1 for parameters). The connectivity architecture is sparse and random with an average connection probability between any pair of cells of 0.2.

The membrane potential V^I^ of each neuron *i* evolves according to the following dynamics^32^:

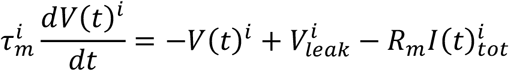

where τ_*m*_ is the membrane time constant, *V*_*leak*_ is the leak membrane potential, *R*_*m*_ is the membrane resistance and ***I***(*t*)_*tot*_ is the total input current. The neuron triggers a spike at time *t*^*^ if *V*(*t*^*^) ≥ *V*_*th****r***_ where *V*_*th****r***_ is the threshold potential. Then, the membrane voltage is set to the reset membrane potential *V*_***r****eset*_for the duration of the absolute refractory time period Δ.

We broke down the term ***I***(*t*)_*tot*_ into two distinct contributions, namely the sum of all the current-based synaptic inputs entering the *i* − *th* neuron ***I***(*t*)_*syn*_ and the external current input ***I***(*t*)_*ext*_, as follows:

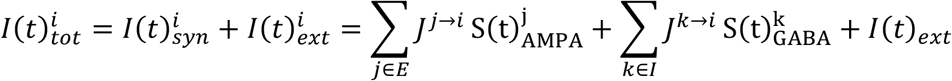

where *J*^*j,k*→*i*^ is the efficacy of the synapsis that connects the *j, k* − *th*pre-synaptic neuron to the *i* −*th* post-synaptic neuron. 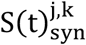 represents the synaptic dynamics which primarily depends on the synapsis type *E*: *AMPA*, ***I***: *GABA*^33^. Every time a *j, k* − *th* presynaptic neuron fires at time *t*^*^, 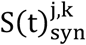 is increased by an amount described by a delayed difference of exponentials, as follows:

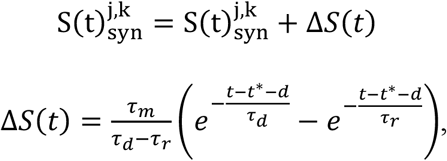

where τ_***r***_ is the rise time, τ_*d*_ is the decay time and *d* is the latency of the synapsis^34^. We defined γ-source activity as the proxy of the local field potential computed by summing the absolute values of the excitatory and inhibitory currents entering the E population^35^.

#### External input into the neural source model

Both populations (excitatory and inhibitory units) receive a time-varying stochastic external current input ***I***(*t*)_*ext*_ that represents the background activity from external (e.g., thalamocortical) afferents, with I neurons receiving more efficacious synapses than E neurons (Supplementary Table 1). The external input was implemented as a series of Poissonian inputs to excitatory synapses with similar kinetics to the recurrent AMPA synapses, but with different efficacy (Supplementary Table 1). These synapses were excited by independent realizations of the same Poissonian process with time-varying input rate *v*_*ext*_(*t*), and therefore, contributing to the single-neuron variability. The *v*_*ext*_(*t*) was composed of the superposition of the signal term *v*_*signal*_(*t*) and the noise term ζ(*t*) as follows:

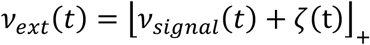

where ⌊·⌋_+_ is the positive part operator.

We modelled the noise component of the input rate ζ(*t*) as a zero-mean Ornstein-Uhlenbeck (OU) process, as follows:

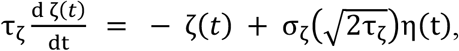

where η(*t*) is the realization of a Gaussian white noise and *σ*_ζ_, τ_ζ_ are the standard deviation and the time constant of the OU process, respectively. τ_ζ_ was set to have a knee in the OU power spectrum at 10 Hz ^36^.

#### Essential dynamics properties of the source activity

We validated essential properties - which have been already extensively investigated elsewhere^34,37^ - of the neural dynamics network with different expressions for *v*_*signal*_(*t*): sustained signals, periodically modulated signals and signals modulated with a Gaussian profile (Supplementary Fig. 2).

First, we modeled external sources with sustained rate with amplitude varying between 2 and 20 spikes/ms. The reverberance of the E-I recurrent connections favored the emergence of reliable broadband γ-oscillations in the [60-150] Hz range (Supplementary Fig. 2). These γ-oscillations encoded the sustained signal increasing the power as the amplitude of the signal increases. It is noteworthy that this frequency range is compatible with the high-gamma speech-related spectral modulation shown in the literature^5,7,38^.

Second, we modeled external inputs with a periodically modulated firing rate, with a varying modulation frequency between 5 and 20 Hz. The source entrained the signal fluctuations generating phase-locked oscillations at the same frequency of the periodic modulation of external input. When a superposition of the sustained and periodic signal was fed into the source, we observed the emergence of two spectral information channels: γ-oscillations that track the mean rate of the signal and low-frequency oscillations entrained by the slow time-scale component in the signal. Interestingly, the phase of the low-frequency oscillations was strongly coupled with the amplitude of γ-oscillations with surges of γ-power close to the peak of the low-frequency oscillation (∼ π/8 phase) (Supplementary Fig. 3)

Finally, to reproduce the temporal pattern of the γ-source activity, e.g., event-locked transient synchronization, we fed gaussian-modulated input rates of external signals in the model (Supplementary Fig. 1,9). The time-activation parameters of the gaussian signal (i.e., fullwidth at half maximum and mean) regulated the duration and temporal focality of the broadband γ-source synchronization (Supplementary Fig. 9).

#### Linear mixing of neural and audio sources

We simulated 99 neural γ-sources and one audio source 𝒮_*a*_ for each simulation. Different sources were fed by different realizations of the *v*_*signal*_(*t*) which expression depends on the simulation scenario (see below and Supplementary Table 1). Therefore, most of the variability at the source-level owes to the stochasticity in *v*_*signal*_(*t*).

In simulation settings we assumed that:

1. One source owes the major contribution to the acoustic-induced artifact (*m* = 1).
2. The audio signal ***z*** = *z*(*t*) used during the denoising pipeline is a good approximation of the artifact source (𝒮_*a*_(*t*) ≈ *z*(*t*)), i.e., the transfer function from audio signal to speech artifact is the identity.
3. The mixing matrix *A* is square.

70% of γ-sources were fed by non-null *v*_*signal*_(*t*) and, as such was considered as “active”. To impose the artifact-to-physiological gamma ratio (AGR), we adjusted the scale of the neural γ-sources by applying a correction factor κ, such that:

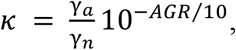

where γ_*a*_ and γ_*n*_ are the power of the audio source and neural sources in the [70-180] Hz frequency range, respectively.

We projected the neural γ-sources and the artifact source into the amplifier space ***X*** applying a random mixing matrix *A*. We included inter-trial variability sampling *A* at each simulation. By imposing the non-singularity of *A*, we avoided ill-posed inversion problems.

Ground truth data ***X***_*gt*_, i.e., noiseless data, were generated by removing the audio source in the mixing operation.

### Simulation scenarios

The *v*_*signal*_(*t*) expression and the procedure used to synthesize the audio source delineated the different simulation scenarios considered in this work, as depicted by Supplementary Fig. 1.

#### Toy examples

We used toy examples for initial assessment of the effect of the PCD pipeline to remove the acoustic-induced artifact in *in-silico* neural signals. We used three scenarios for these toy examples, where *v*_*signal*_(*t*) was defined as the superposition of the sustained and periodically modulated signal, as follows:

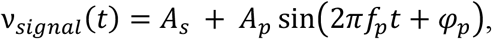

where *A*_*s*_ is the amplitude of the sustained input, and *A*_*p*_, *f*_*p*_ and *φ*_*p*_ are the amplitude, the frequency and the phase of the periodic signal, respectively.

In Scenario 1 (Sinusoidal audio scenario, SAS), we synthetized the contaminating audio source 𝒮_*a*_ as a sinusoidal function with noise, as follows:

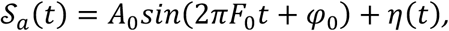

where *A*_0_ is the amplitude, *F0* is the fundamental frequency, *φ*_0_ is the phase, and η(*t*) is the white noise term.

In Scenario 2 (Colored noise audio scenario, CAS), the audio source was synthesized by coloring the spectrum of the white noise η(*t*), as follows:

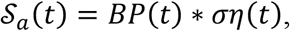

where BP(t) is the impulse response of the 25^th^ order Butterworth filter, *σ* is the standard deviation and * is the convolution operator.

In Scenario 3 (Modulated colored noise scenario, MCAS), we mimicked the temporal profile of the audio in the Syllable Repetition Task (see *Triplet repetition task data*), applying the activation pattern *M*(*t*) to the audio source in Scenario 2, which reads:

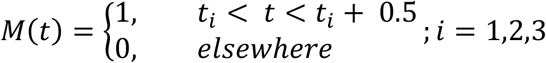

where *t*_*i*_ is the onset of *i* − *th* syllable, which lasts 0.5 s.

#### Realistic scenario

In this Scenario (recorded audio scenario, RAS), we simulated 60 trials of speech-locked transient γ-source activity, feeding Gaussian signals in the network, as follows:

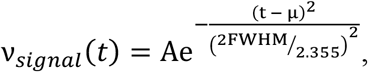

where A and μ are the intensity and temporal location modulation, and *FWHM* is the fullwidth at half maximum which controls the temporal focality of the modulation. Parameters were set to reproduce physiological speech-locked γ-modulation patterns described in literature^4,39,40^. As audio source, we selected the recorded audio signals during the Syllable Repetition Task with different audio spectrogram profiles, e.g., non-stationary pitch (Supplementary Fig. 10).

#### Computational simulation settings

The simulations were performed using a finite difference integration scheme based on the 2nd-order Runge-Kutta algorithm with discretization time step Δ*t* = 0.05 *ms*. to update the neural dynamics. Simulation duration varied between 1 s and 10 s. We did not include the first 200 ms of each simulation in the analysis to avoid transitory dynamics in the network. We fixed the seeds of the random generator number for sake of the reproducibility across different simulation sessions.

#### Triplet repetition task data

54 English-speaking patients undergoing an awake DBS implantation surgery consented to participate in an intraoperative speech task at the University of Pittsburgh (IRB Protocol #PRO13110420). Patients were instructed to repeat consonant-vowel syllable triplets that were played on their earphones. Specifically, the basis set of phonemes included 4 consonants (/v/, /t/, /s/,/g/) and 3 cardinal vowels (/i/, /a, /u/) with distinctive tongue positions and acoustic properties. 1 to 4 recording sessions of up to 120 syllable triplets each were performed by participants. Sessions differed regarding the neural recording modalities registered. Electrocorticography (ECoG) was always registered through one or two strips of 53 or 64 contacts targeted to cover the left ventral sensorimotor cortex, superior temporal gyrus and inferior frontal regions. MER and LFPs from a macro ring located 3mm above the microelectrode tip were registered during subcortical target mapping, for participants undergoing subthalamic nucleus (STN) or Globus Pallidus Internus (GPi) DBS. LFP registered from the DBS lead were recorded from the STN, GPi and ventral intermediate nucleus (VIM). The produced audio signal was recorded by a directional microphone placed near the patient’s mouth. Data was time-aligned, pre-processed, low-pass filtered at 500Hz, down-sampled to 1kHz and saved as a FieldTrip^41^ object for subsequent analyses. For more information about the dataset, we refer the reader to Bush et al. 2021^8^.

### Algorithm implementation

#### Trial-wise denoising

Since phase relationships between the artifact source and the neural recordings is not consistent across trials (Supplementary Fig. 12), the PCD pipeline was applied on a trial-wise basis. Data was high-pass filtered above 2 Hz and notch-filtered at 60 Hz and its three first harmonics. Given that the artifact was observed only during overt speech production times, epochs around the produced speech onset were extracted for fitting the model. Under the hypothesis that the artifact is introduced in the acquisition chain, the different synchronized brain recording modalities (MER-LFP + ECoG or DBS-LFP + ECoG) were combined to form a unique data matrix. That is, the signals from all available brain recording modalities were treated as a unique sensor space.

For every trial, the model was fitted as follows. First, the epoched audio signal was used to estimate the SAFB, and thus the noise and signal band were accordingly defined (Box 1 - step 1). Data preparation proceeded, which included applying the Hilbert transform to find the analytic representation of the signal, as well as data whitening. The audio signal was z-scored for normalization purposes and used as the best guess of the artifact source. The SSD algorithm was applied to the real part of the signal. The final number of SSD components to keep (*k*) was automatically selected based on the participation ratio (PR), defined as follows:

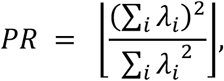

where 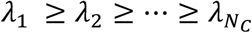 are the eigenvalues resulting from solving SSD which account for the SNR participation of each SSD component. The PR has been shown to be a reasonable measure of neural dimensionality in PCA^42^. PCO was applied on data projected onto the SSD space, extracting one component at the time. Every component resulted in a mean vector length (MVL) value, such that MVL_1_ ≥ MVL_2_ ≥ … ≥ MVL_*k*_. Once both spatial filtering methods were learned, the unified unmixing and mixing matrix that described the PCD pipeline were computed (Box 1 – step 4).

The identification of the artifactual sources was also automatically done by finding the elbow in the trace of the MVL value across components, that is the point at maximum curvature differentiating the contribution of strong and weak components. Those components showing the highest MVL values were identified as artifactual.

Considering that further analysis would be done on the data, a wider epoch starting 6 seconds before and ending 6 seconds after the produced onset was used when applying the learned matrix transformation.

#### Algorithm stress test on simulated data: parametric sweep and performance evaluation

We applied the PCD pipeline on simulated data using the speech artifact 𝒮_*a*_(*t*) as audio signal (𝒮_*a*_(*t*) ≈ *z*(*t*)). Because we simulated only one artifactual source, we expected the PCO to find only one component with high coherence with the audio (i.e. MVL) in the PCO space. To assess the performance of the denoising pipeline, we calculated time-, frequency- and phase-domain metrics that estimate the agreement between the ground-truth ***X***_***gt***_ and cleaned data ***X***, and *z*(*t*) and *z*(*t*)_*est*_.

For each channel, we compared the similarity in the time-domain of the neural data (***X*** *νs*· ***X***_***gt***_), as follows:

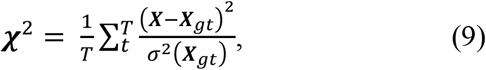

where *T* is the duration of the simulation and *σ*^2^(·) is the variance. ***χ*** values have been converted to a logarithmic scale for visualization purposes (Supplementary Fig. 5,7).

To assess the fidelity of the estimated speech-induced artifact 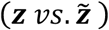, we used the magnitude coherence estimate (MSCE) and the consistency of the phase difference based on the PLV. The MSCE returns values between 0 and 1 indicating how similar two signals are at each frequency, as follows:

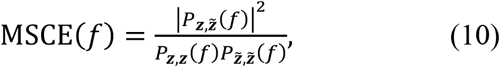

where *P* stands for the power spectral density. The PLV is a metric between 0 and 1 that quantifies the phase agreement between two signals, as follows:

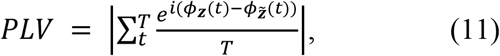

where *ϕ* is the instantaneous phase.

Moreover, to stress the PCD pipeline in the different scenarios, we evaluated performance during a sweep of key parameters, including the AGR [-100, 30] dB, *F0* [70, 180] Hz and *N*_*c*_ [3, 100] in the SAS scenario. We swept Δ*F* [2, 16] *Hz*, the filter order [3, 27] and the simulation duration [0.5, 3.5] s in the CAS scenario.

Results showed that for pure sinusoidal artifacts, PCD perfectly removed the artifact regardless of the AGR, fundamental frequency and number of channels (Supplementary Fig. 4,5). For the colored noise artifact, simulations suggested a small decrease of performance with broadband artifacts, which is consistent with known limitations of SSD (Supplementary Fig. 6,7). Finally, PCD yielded robust performances when tested with modulated colored noise artifact simulations (Supplementary Fig. 8).

#### Neural preservation assessment

Principal Component Analysis (PCA) is a dimensionality reduction technique that identifies an ordered set of orthogonal directions that captures the greatest variance in the data. It is widely accepted and used in the neuroscience community for analyzing neural population activity^43^. The low-dimensional space identified by PCA captures variance of all types, including noise. Such data representation can be thought of as the Cartesian coordinate basis describing subspaces in which the data lie^44^. Thus, for assessing neural preservation after applying denoising methods, the PCA low-dimensional space was utilized. Subspaces describing the same geometry as the ground truth data should be found after denoising if neural preservation is achieved. Thus, for every denoising pipeline, as well as the ground truth and the noisy data, PCA was fitted independently. The learned loading and scores were plotted for each decomposition made on the first 3 principal components (PC).

To quantify the degree of similarity between the PCA loadings 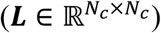of the ground-truth data and the resulting data after each denoising pipeline was applied, the cosine similarity (CS)^45^ was used. CS measures the extent to which two vectors point in the same direction. Given that PCA loading signs are arbitrary, bounded CS between 0 and 1 can be found by taking the absolute value of the cosine similarity, as follows:

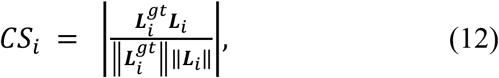

where *i* denotes the index of a given loading vector, ***L***_***gt***_ indicates the PCA loading matrix of the ground-truth data ***X***_***gt***_, ***L*** stands for the PCA loading matrix of a given denoised data ***X***_*denoised*_, and ‖·‖ and |·| denotes the ℓ_2_ − norm and the absolute value operators, respectively. The closer CS is to 1, the more similar the two vectors are. Thus, a good denoising pipeline from the neural data preservation point of view should be the one from which PCA loading vectors resemble the same directions (see Fig. 4d).

#### Inter-trial coherence for speech artifact quantification

The speech artifact level at each electrode was computed based on the inter-trial phase consistency (ITPC)^46^, following the same framework proposed and used in Bush et al. 2021. That is, the audio and the neural signals were band-pass filtered between 70 and 240 Hz, i.e., within the plausible SAFB range. Then considering the complex representation of the neural data for a given channel ***y*** = ***x*** + *i*ℋ(***x***) and the audio signal ***z*** at a given trial *e*, the phase between these two quantifies can be measured by:

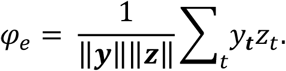

At the end of this procedure, all the phases across the *N*_*t*_ trials are arranged on a vector 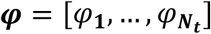. If there is inter-trial phase consistency, the mean value of ***φ*** across trials (*⟨****φ****⟩*) will be different from 0, and thus it can be quantified as follows:

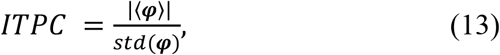

where *std*(·) stands for the stand deviation. It has been found that ITPC values equal or above 3.08^8^ indicates that the presence of the speech artifact on that given electrode is significant, and thus the electrode must be considered contaminated^8^.

#### Artifact presence quantification: definition of homogeneity, strength and clean electrode gain

Artifact homogeneity should quantify the consistency of the artifact presence across electrodes. Let us denote 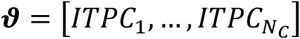 as a vector with the stored ITPC value found per each electrode.

Considering that *n* −dimenstional unit vectors have variance between 0 and 1/*n*, following the idea proposed by Umakantha et al.^47^ we define the artifact homogeneity as follows:

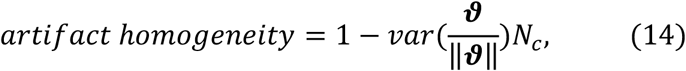

that is, a value between 0 and 1, for homogeneous artifact presence and non-homogeneous artifact presence, respectively. Artifact strength was directly measure by the mean ITPC value found across electrodes:

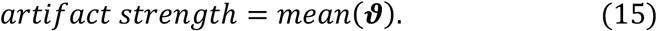

The clean electrode gain was computed as the relative change of the clean electrode percentage before (%*CE*_*denoised*_) and after (%*CE*_***r****aw*_) applying a denoising method, that is:

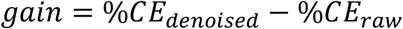

#### Deep learning model for consonant identification

Densely-connected convolutional neural networks (DenseNet)^24^ were trained to classify the consonants from neural signals (ECoG). For each syllable, original/denoised ECoG signals were spectrally filtered into 7 frequency bands ranging between 0 to 250 Hz. These syllable-level neural data were used as training set. We then extracted the perceptual linear prediction (PLP)^48^ features from the corresponding audio recordings. Both PLP features and consonant identities were used as training labels. Our DenseNet model was designed to first map neural signals into PLP spectra and then predict the consonant class from the PLP space. For that purpose, the mean-squared-error loss of PLP feature prediction and the cross-entropy loss of consonant classification were jointly optimized during model training. We used 5-fold cross-validation while measuring model performance, withholding a different 20% partition from the training data as test set each time. Separate models were trained for each subject and each data type, resulting in 12 different models (3 subjects, 4 data types: Raw, CAR, ICA, PCD). During testing, reserved ECoG data were fed into the trained model, and the accuracy was measured based on the consonant predictions.

### Traditional methods for denoising

#### CAR as a spatial filtering algorithm

Common average reference (CAR) is a spatial filtering method that subtracts the common noise at every electrode, calculated as the average of all recordings. The data re-referenced via CAR is calculated as follows:

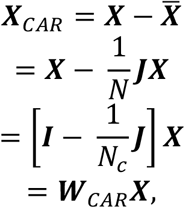

where *N*_*c*_ accounts for the number of electrodes and 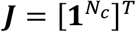 is an *N*_*c*_ × *N*_*c*_ matrix of ones. With this formulation it is easy to see that CAR can be thought of as a spatial filtering method which implies a linear transformation of the data.

Given that CAR takes the average across channels, data structure is not critical in this matter. The matrix transformation was applied in the continuous data for each type of channel recording.

#### The PCA + ICA pipeline

Independent component analysis (ICA) assumes that the sources linearly mixed in (1) are independent. Such an assumption is true for many artifacts that appear in brain recordings, like electrocardiography or electromyography. In order to ensure that independent sources are extracted from the data, non-gaussianity of ***W***^*T*^***X*** is maximized. Here in particular, ICA is implemented using Picard^49^, a fast algorithm to solve the maximum likelihood estimation ICA formulation. This ICA implementation was chosen since it is known to result in robust components estimations in cases where the sources are not completely independent.

Basic data pre-processing was done before applying ICA. A temporal filtering above 2 Hz was implemented in order to remove low-frequency drifts, which can negatively affect the quality of the ICA fit^50^. Then data was z-scored and PCA was applied in order to both feed ICA with whitened data and to reduce the dimensionality of the problem. The number of PCA components was automatically selected by setting the explained variance to 0.99. As in the PCD pipeline, model fitting was done using the windowed data within the produced audio epoch, combining, if exists, the different brain recording modalities. For the sake of comparison with PCD, artifactual sources identification was done based on the circular mean of the phase difference. Those components showing the highest phase-locking values were selected as artifactual. Denoising via low-rank factorization was applied in the wider epochs, as done for PCD.

### Computing environment

Data simulation and numerical experiments were conducted in Matlab 2020b. We used custom-based scripts (BML toolbox; https://github.com/Brain-Modulation-Lab/bmlbmlbmlbmlbml based on the Fieldtrip library (https://www.fieldtriptoolbox.org). We used online available implementations of SSD (https://github.com/svendaehne/matlab_SSD) and PCO (https://github.com/neurophysics/PCO_matlab). For measuring the phase-locking values and phase differences the CircStast toolbox (https://www.jstatsoft.org/article/view/v031i10) was utilized. In addition, Rstudio was used to compute the ITPC. For computing the simulations, we used the publicly available C-optimized implementation (http://senselab.med.yale.edu/ModelDB/ShowModel.asp?model=152539)^33^. We computed the MSCE and the consistency of the phase difference by using the built-in MATLAB *mscohere* function and *circ_r* function in the CircStat Toolbox^51^, respectively. We used the RainCloud library^52^ to compare distributions of data (https://github.com/RainCloudPlots/RainCloudPlots#read-the-preprint).

## Data availability

The data that support the findings of this study are available upon reasonable request. A formal data sharing agreement is required.

## Code availability

The code for the PSD algorithm is available online at https://github.com/Brain-Modulation-Lab/PCD

## Acknowledgements

We would like to thank research participants for their generous contribution of time and effort in the operating room and additional experimenters who acquired and organized the data. This work was funded by the National Institute of Health (BRAIN Initiative), through grants U01NS098969, U01NS117836 and R01NS110424 to R.M.R.

## Author contributions

V.P., M.V. and A.B. conceived the computational experiments, and R.M.R conceived the experimental task design. V.P conceived and coded the initial algorithm. V.P and M.V designed the final algorithm and pipeline. A.B performed the data curation. M.V. designed and coded the simulated data framework. V.P. and M.V. tested the algorithm against real and simulated data. S.L. and Q.R. did the deep-learning experiments. V.P, M.V, S.L, Q.R. and A.B contributed tools to curate and analyze data. A.B., N.E.C and R.M.R gave insights into analyzing results. The original draft was written by V.P and M.V. and was revised and edited by all authors. R.M.R supervised the project.

