## Supplementary for "A supervised data-driven spatial filter denoising method for speech artifacts in intracranial electrophysiological recordings"

### Supplementary Tables

**Table S1: LIF network parameters and external input definitions for different in-silico scenarios.** List of parameters used for in-silico brain sources. $\mathrm{N}(\mu, \sigma)$ indicates the gaussian distribution with mean μ and standard deviation $\sigma$. $U(a, b)$ defines the continuous uniform distribution with support $[a, b]$.

| | | |
|---|---|---|
| **Network structure** | | |
| Number of neurons | $N_E$ | 4000 |
| | $N_I$ | 1000 |
| Connection probability | $p$ | 0.2 |
| **Neuron dynamics** | | |
| Discretization step | $\Delta t$ | 0.05 ms |
| Leak membrane potential | $V_{leak}^{E}$ | -70 mV |
| | $V_{leak}^{I}$ | -70 mV |
| Threshold membrane potential | $V_{thr}^{E}$ | -52 mV |
| | $V_{thr}^{I}$ | -52 mV |
| Reset membrane potential | $V_{reset}^{E}$ | -59 mV |
| | $V_{reset}^{I}$ | -59 mV |
| Absolute refractory period | $\Delta^{E}$ | 2 ms |
| | $\Delta^{I}$ | 1 ms |
| Membrane resistance | $R_m^E$ | 0.04 $G\Omega$ |
| | $R_m^I$ | 0.05 $G\Omega$ |
| Membrane time constant | $\tau_m^E$ | 20 ms |
| | $\tau_m^I$ | 10 ms |
| **Synapses** | | |
| Synaptic efficacy | $J^{E\to E}$ | -10.5 pA |
| | $J^{I\to E}$ | 42.5 pA |
| | $J^{ext\to E}$ | -13.75 pA |
| | $J^{I\to I}$ | 54 pA |
| | $J^{E\to I}$ | -14 pA |
| | $J^{ext\to I}$ | -19 pA |
| Synaptic delay | $d^{E\to E}$ | 1 ms |
| | $d^{I\to E}$ | 1 ms |
| | $d^{ext\to E}$ | 1 ms |
| | $d^{I\to I}$ | 1 ms |
| | $d^{E\to I}$ | 1 ms |
| | $d^{ext\to I}$ | 1 ms |
| Synaptic rise constant | $\tau_r^{E\to E}$ | 0.40 ms |
| | $\tau_r^{I\to E}$ | 0.25 ms |
| | $\tau_r^{ext\to E}$ | 0.40 ms |
| | $\tau_r^{I\to I}$ | 0.25 ms |
| | $\tau_r^{E\to I}$ | 0.20 ms |
| | $\tau_r^{ext\to I}$ | 0.20 ms |
| Synaptic decay constant | $\tau_d^{E\to E}$ | 2 ms |
| | $\tau_d^{I\to E}$ | 5 ms |
| | $\tau_d^{ext\to E}$ | 2 ms |
| | $\tau_d^{I\to I}$ | 5 ms |

| | | |
|---|---|---|
| | $\tau_d^{E\to I}$ | 1 ms |
| | $\tau_d^{ext\to I}$ | 1 ms |
| **External input $I_{ext}(t)$ in Toy examples: SAS, CAS, MCAS** $A_s + A_p \sin\left(2\pi f_p t + \varphi_p\right) + \zeta(t)$ | | |
| Static amplitude | $A_s$ | $U(12,16)$ $spike/(ms * cell)$ |
| Periodic amplitude | $A_p$ | $N\left(0, \frac{16}{3}\right) spike /(ms * cell)$ |
| Frequency oscillation | $f_p$ | $N(10,1.5)$ Hz |
| Phase oscillation | $\varphi_p$ | $U(-\pi, \pi)$ |
| UO constant time | $\tau_\zeta$ | 0.16 s |
| Uo standard deviation | $\sigma_\zeta$ | 4 spikes/ms |
| **External input $I_{ext}(t)$ in Realistic simulations: RAS** $\mathrm{A}e^{-\frac{(\mathrm{t}-\mu)^2}{\left(2\mathrm{FWHM}/2.355\right)^2}} + \zeta(\mathrm{t})$ | | |
| Amplitude modulation | $A$ | $U(10,14)$ $spike/(ms * cell)$ |
| Peak modulation time | $\mu$ | $N(5,\ 0.05)$s |
| Full width at half maximum | $FWHM$ | $N(1.2,\ 0.15)$s |
| UO constant time | $\tau_\xi$ | 0.16 s |
| Uo standard deviation | $\sigma_\zeta$ | 4 spikes/ms |

**Table S2: Audio definitions for the different toy examples: SAS, CAS, MCAS.** List of parameters used for in-silico audio sources. $\sigma\eta(t)$ indicates the white noise with mean 0 and standard deviation $\sigma$.

| | | | |
|---|---|---|---|
| **Toy examples** | **Sinusoidal audio scenario (SAS):** $A_0 sin(2\pi F_0 t + \varphi_0) + \eta(t)$ | | |
| | Amplitude audio source | $A_0$ | $15 \cdot 10^3$ |
| | Fundamental frequency | $F_0$ | 120 Hz |
| | Phase audio source | $\varphi_0$ | $\pi/3$ |
| | **Colored noise audio scenario (CAS):** $\sigma\eta(t)$ | | |
| | Standard deviation white noise | $\sigma$ | $15 \cdot 10^{15}$ |
| | Center frequency filter | $F_0$ | 120 Hz |
| | Bandwidth filter | $\Delta F$ | 5 Hz |
| | Order filter | $b$ | 25 |
| | **Modulated colored noise audio scenarios (MCAS):** $\sigma\eta(t)$ → BP filter → $M(t)$ → | | |
| | Standard deviation white noise | $\sigma$ | $15 \cdot 10^{15}$ |
| | Center frequency filter | $F_0$ | 120 Hz |
| | Bandwidth filter | $\Delta F$ | 5 Hz |
| | Order filter | $b$ | 25 |
| | # bumps | | 3 |
| | Duration bump | | 0.5 s |

### Supplementary Figures

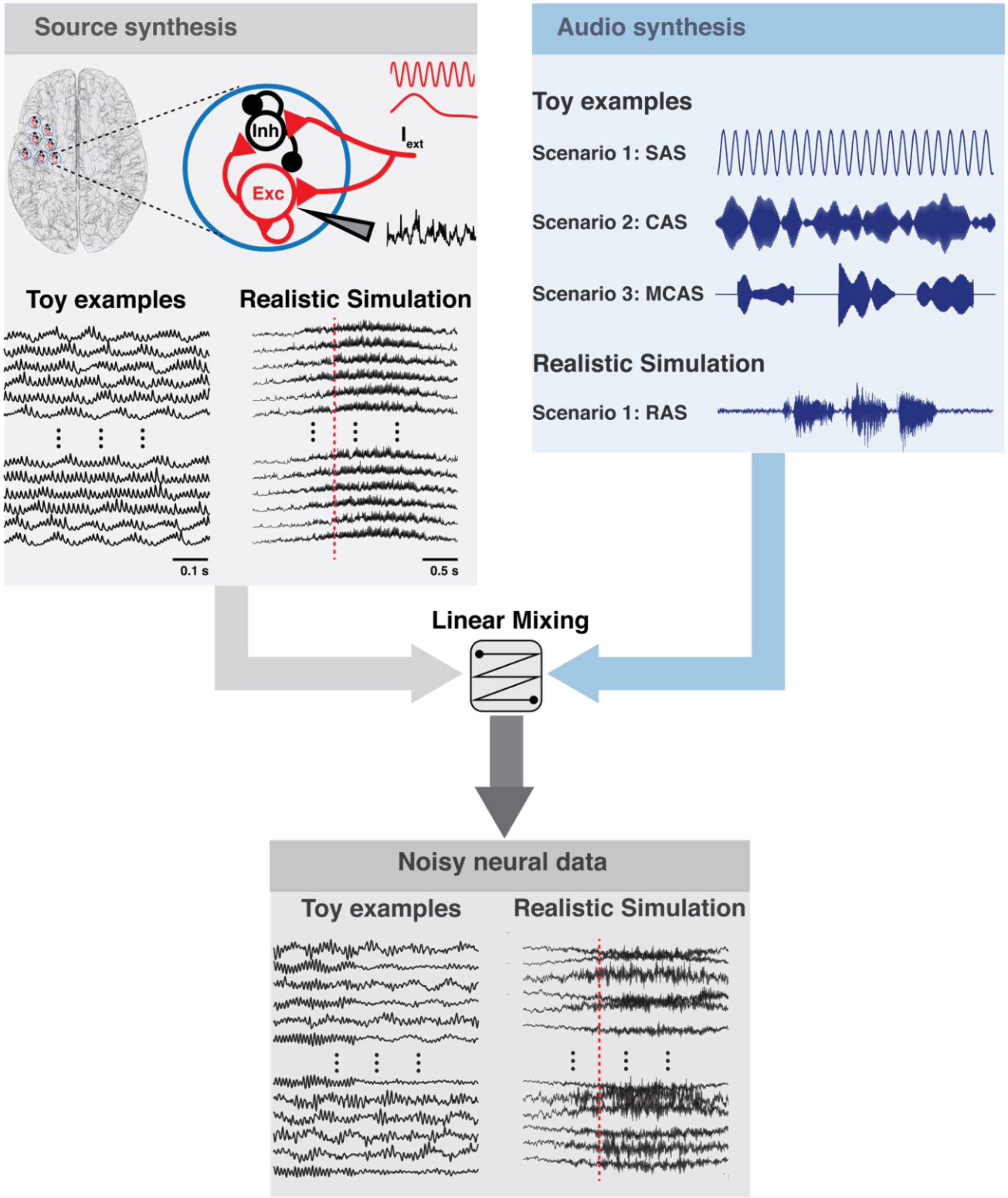


**Supplementary Fig. 1 | Schematic of the pipeline for the simulation scenarios.** Brain sources are simulated using a sparse (sparseness; p = 0.2) LIF network of excitatory (N= 4000, red line) and inhibitory neurons (N = 1000, black line). Both populations receive recurrent activity and external excitatory inputs $I_{ext}$. $I_{ext}$ is a Poissonian process with time-varying input rate $\nu_{ext}(t)$. The size of the synaptic connection (inhibitory: circle and excitatory: triangle) depicts the synaptic efficacy. LFPs are estimated using a simple computational proxy which neglects the direct contribution of the inhibitory population (refer to 1 for details). Simulation of neural data affected by the vibration artifact are obtained by linearly mixing brain sources and audio signal by the application of a mixing matrix. We simulated different scenarios according to the type of audio signal and the $\nu_{ext}(t)$ expression (refer to Methods for details). The vertical red dashed line depicts the onset of the speech production event. (SAS: sinusoidal audio scenario, CAS: colored noise audio scenario, MCAS: modulated colored noise scenario, RAS: recorded audio scenario)

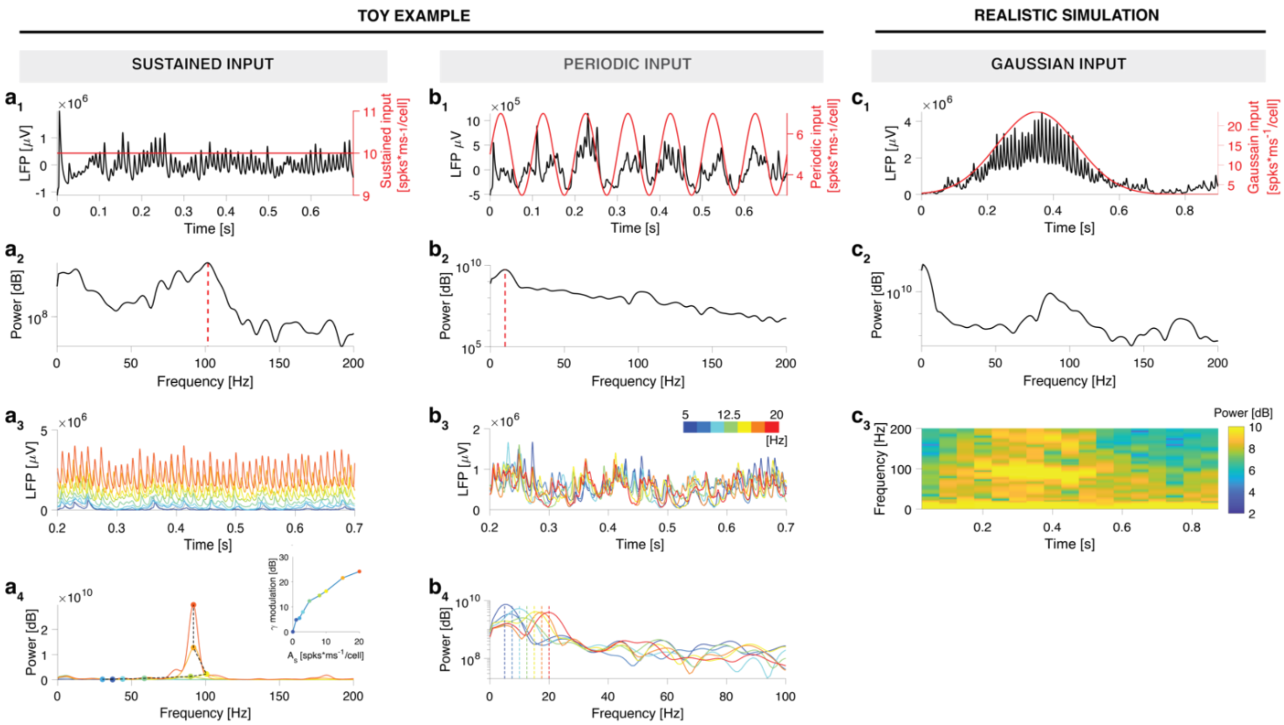


**Supplementary Fig. 2 | Source activity reflects temporal properties of the network input $\nu_{signal}(t)$. $a_1$- $a_2$**, Temporal and spectral source activity during the injection of a sustained input at 10 spikes/(ms*cell) (red line). Strong $\gamma$-oscillations are visible in the time domain with a peak at ~100 Hz (vertical red dashed line). **$a_3$- $a_4$**, gamma-oscillations entrainment can be tweaked by sweeping the intensity of the sustained input, as depicted by the color code. **$b_1$- $b_2$**, Temporal and spectral source activity during the injection of a periodically modulated input rate at 10 Hz (red line). Source activity tracks the frequency of the periodic input, revealing a peak at 10 Hz (vertical red dashed line). **$b_3$- $b_4$**, High fidelity between the frequency of the periodic input (vertical dashed line) and the peak of the oscillatory source activity, as depicted by the color code. **$c_1$- $c_3$** Time-frequency representation of the source activity during the injection of a gaussian-modulated input. Transient gamma-oscillations emerge between 0.2 s and 0.5 s.

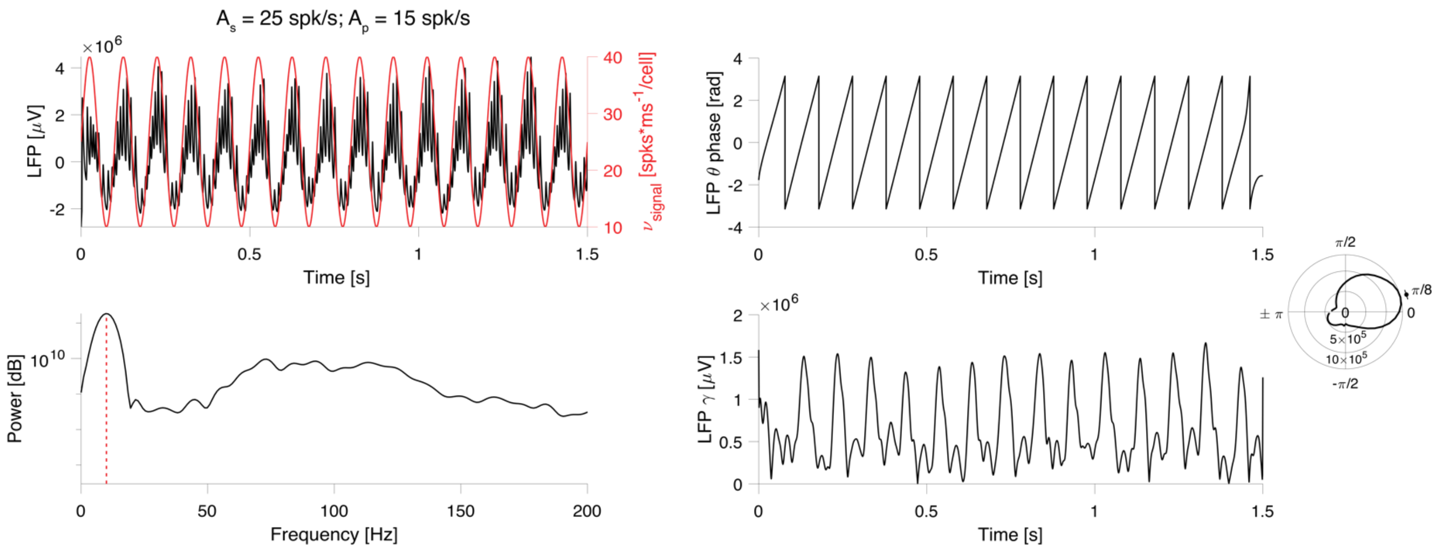


**Supplementary Fig. 3 | Neural source activity reproduces physiological phase-amplitude coupling.**
Temporal and spectral source activity during the injection of a periodic input at 10 Hz (red line). Source activity tracks the oscillation frequency of the period input (vertical red dashed line). When dissecting spectral components of the source activity, phase of the low-frequency oscillations was strongly coupled with the amplitude of $\gamma$-oscillations with surges of $\gamma$-power close to the peak of the low-frequency oscillation ($\sim \pi/8$ phase). Inset plot shows $\gamma$-power w.r.t the phase of the low frequency oscillation.

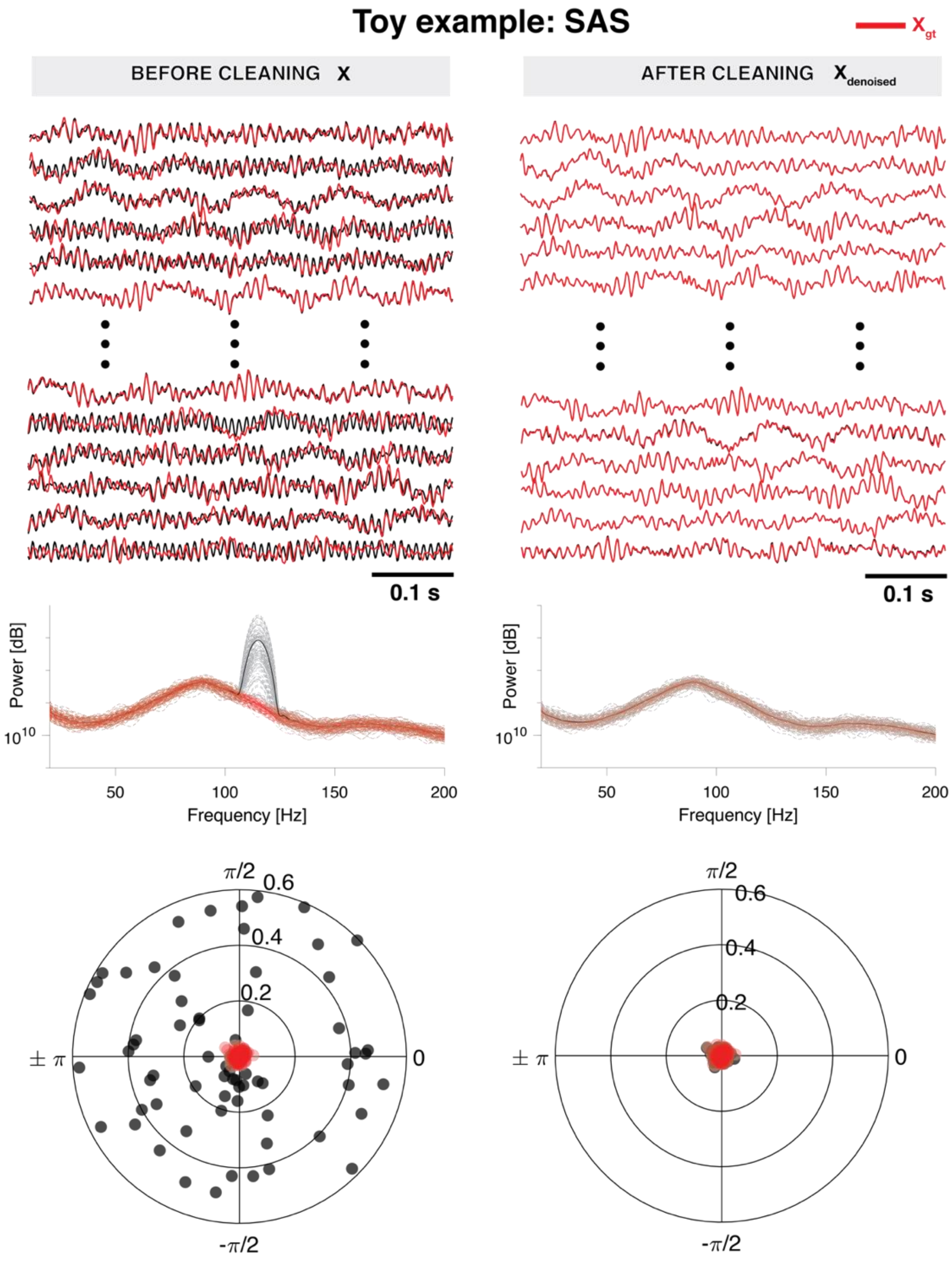


**Supplementary Fig. 4 | PCD perfectly removes the audio source in the toy example SAS.**
Noisy neural sources (**$X$**, black line, left) are obtained by linear mixing of ground-truth neural sources ($\boldsymbol{X_{gt}}$, red line) with a sinusoidal audio source ($F_0 = 120\ Hz$). **$X$** exhibits a narrowband component around the fundamental frequency $F_0$ as well as phase locking with the audio source. After cleaning, $\boldsymbol{X_{denoised}}$ (black line, right) resembles $\boldsymbol{X_{gt}}$ in the time and frequency domain. Finally, $\boldsymbol{X_{denoised}}$ is uncoupled with the audio source.

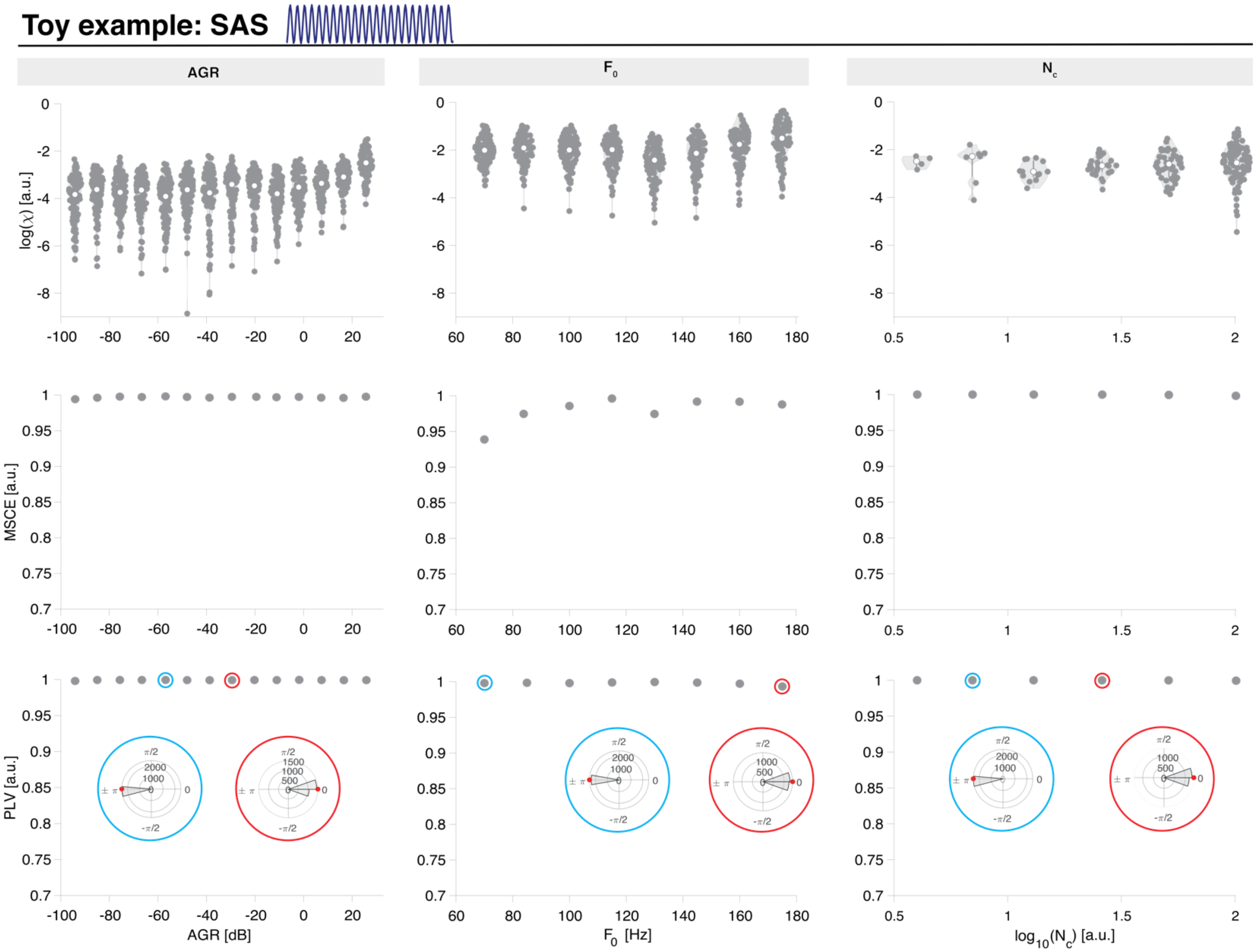


**Supplementary Fig. 5 | Performances of PCD pipeline in the toy example SAS across a range of AGR, $\boldsymbol{F_0}$, and $\boldsymbol{N_C}$.**
PCD pipeline is robust to AGR (**a**), $\boldsymbol{F_0}$ (**b**) and $\boldsymbol{N_C}$ (**c**) changes in terms of agreement between the ground-truth $\boldsymbol{X_{gt}}$ and cleaned data $\boldsymbol{X}$ (top, $\log(\chi)$) and between the artefact z(t) and the estimated artefact $z(t)_{est}$ (center-bottom, MSCE and PLV). Inset polar plots show the phase difference (z(t) vs. $z(t)_{est}$) distribution in two exemplary simulations (red: correlation, cyan: anticorrelation), as pinpointed by surrounding circles Red filled circle displays the average.

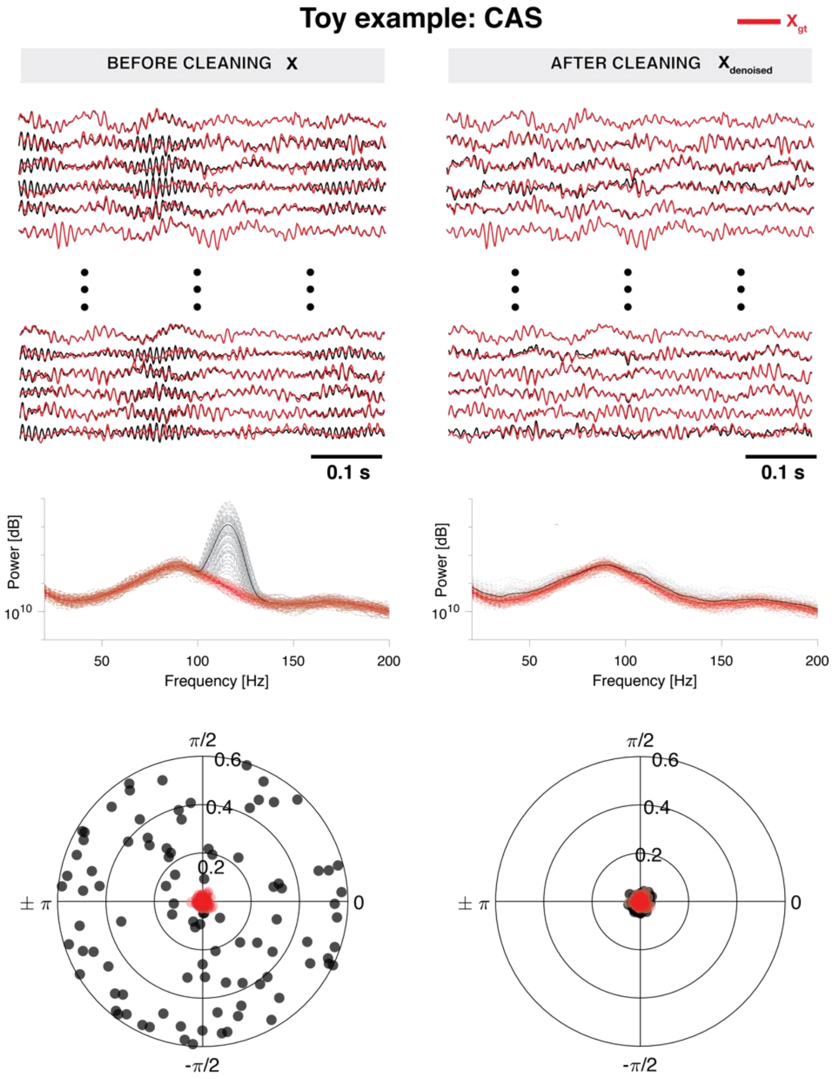


**Supplementary Fig. 6 | PCD perfectly removes the audio source in the toy example CAS.** 60 trials of noisy neural sources **(**$\boldsymbol{X}$, black line, left) are obtained by linear mixing of ground-truth neural sources ($\boldsymbol{X_{gt}}$, red line) with a colored noise audio source ($F_0 = 120\ Hz, \Delta F = 5\ Hz$, duration bump = 0.5 s).$\boldsymbol{X}$ exhibits a narrowband component around the fundamental frequency $F_0$ as well as phase locking with the audio source. After cleaning, $\boldsymbol{X_{denoised}}$ (black line, right) resembles $\boldsymbol{X_{gt}}$ in the time and frequency domain. Finally, $\boldsymbol{X_{denoised}}$ is uncoupled with the audio source. Performance metrics ($\log(\chi)$, MSCE and PLV) distributions are displayed.

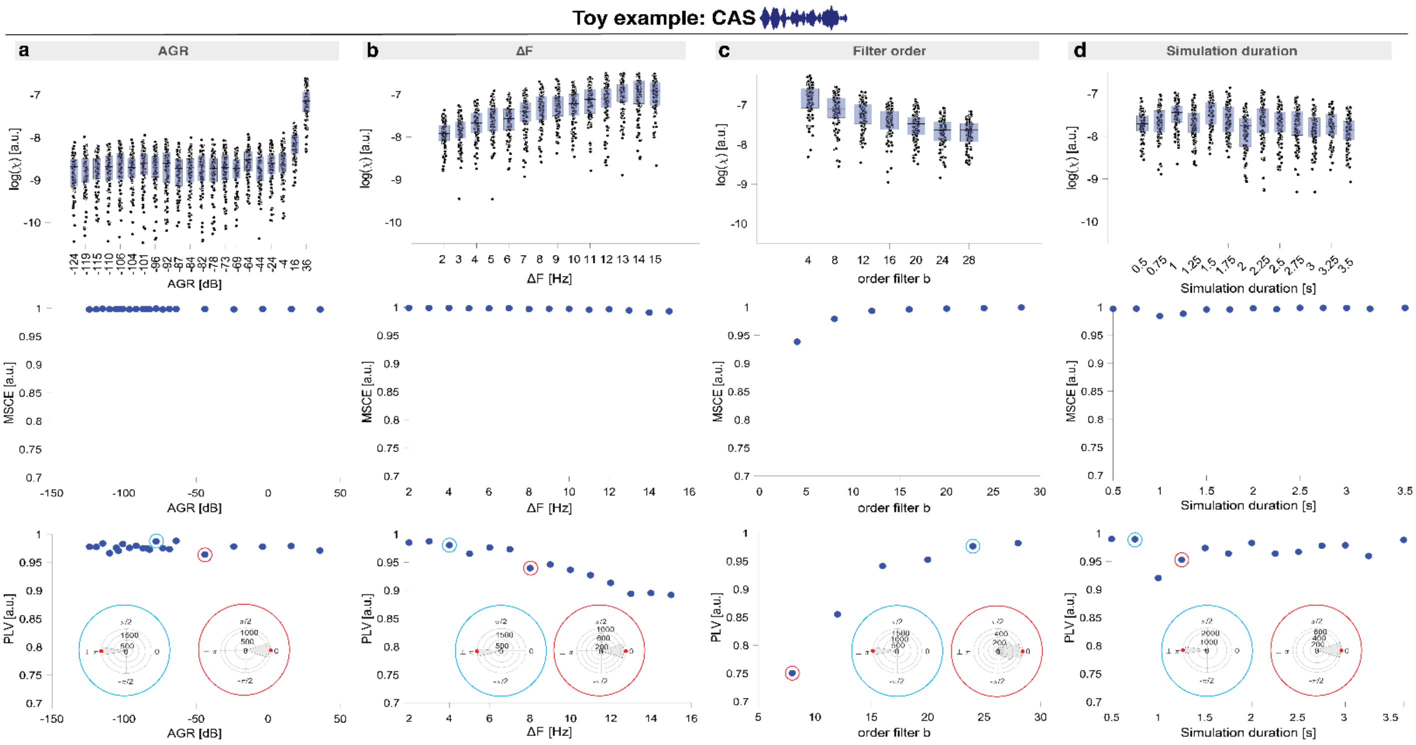


**Supplementary Fig. 7 | Performances of PCD pipeline in the toy example CAS across a range of AGR, ΔF, filter order b and simulation duration. a**, PCD pipeline is robust to artifact-to-physiological gamma ratio (AGR) changes in terms of agreement between the ground-truth $\boldsymbol{X}_{gt}$ and cleaned data $\boldsymbol{X}$ (top, $log(\chi)$) and between the artifact $z(t)$ and the estimated artifact $z(t)_{est}$ (center-bottom, MSCE and PLV). **b-c, Broad SAFB significantly reduces PCD performances.** PCD pipeline is more accurate to remove narrowband artifacts, as suggested by the drop in performances when either the artifact frequency peak is too large (ΔF) or not well defined (order filter b) (Methods, CAS artifact definition). **d,** Duration of the simulation does not significantly impact PCD performances. Inset polar plots show the phase difference ($z(t)$ vs. $z(t)_{est}$) distribution in two exemplary simulations (red: correlation, cyan: anticorrelation), as pinpointed by surrounding circles Red filled circle displays the average.

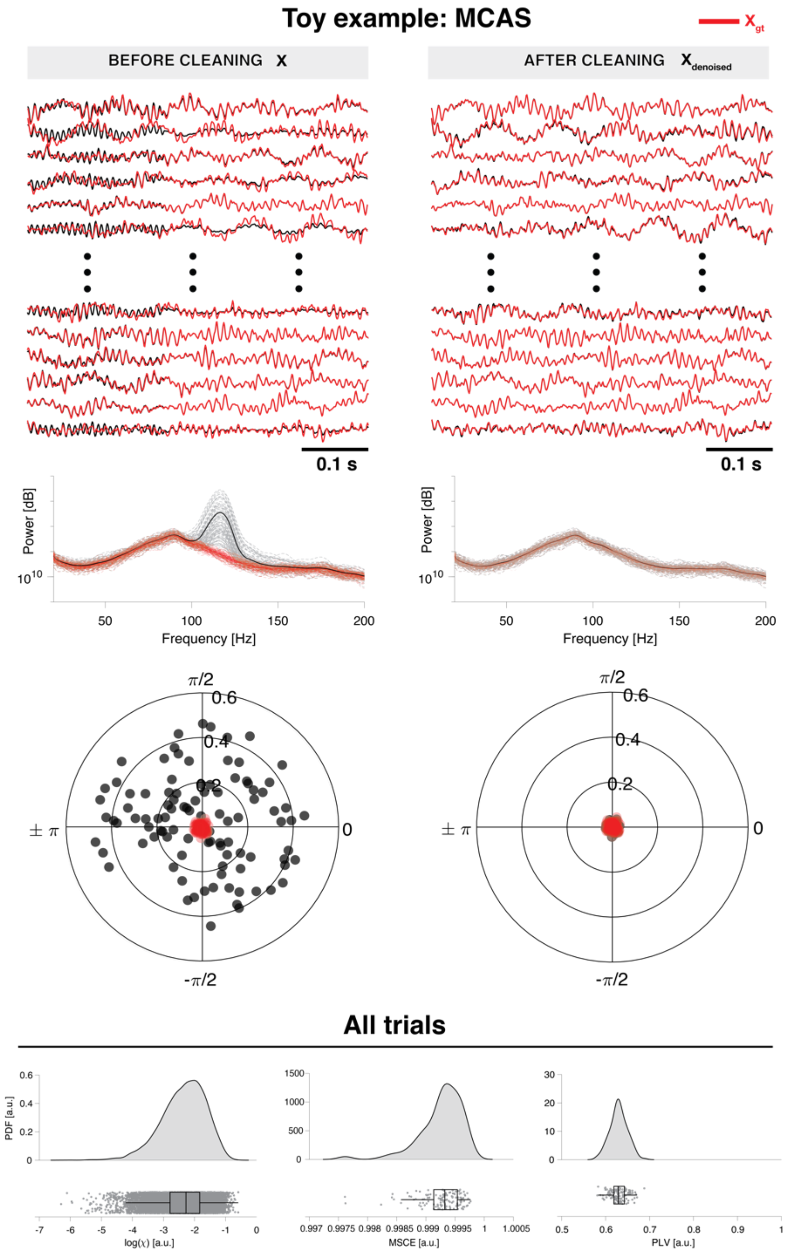


**Supplementary Fig. 8 | PCD perfectly removes the audio source in the toy example MCAS.**
Noisy neural sources (**$\boldsymbol{X}$**, black line, left) are obtained by linear mixing of ground-truth neural sources ($\boldsymbol{X_{gt}}$, red line) with a modulated colored noise audio source ($F_0 = 120\ Hz, \Delta F = 5\ Hz$). $\boldsymbol{X}$ exhibits a narrowband component around the fundamental frequency $F_0$ as well as phase locking with the audio source. After cleaning, $\boldsymbol{X_{denoised}}$ (black line, right) resembles $\boldsymbol{X_{gt}}$ in the time and frequency domain. Finally, $\boldsymbol{X_{denoised}}$ is uncoupled with the audio source.

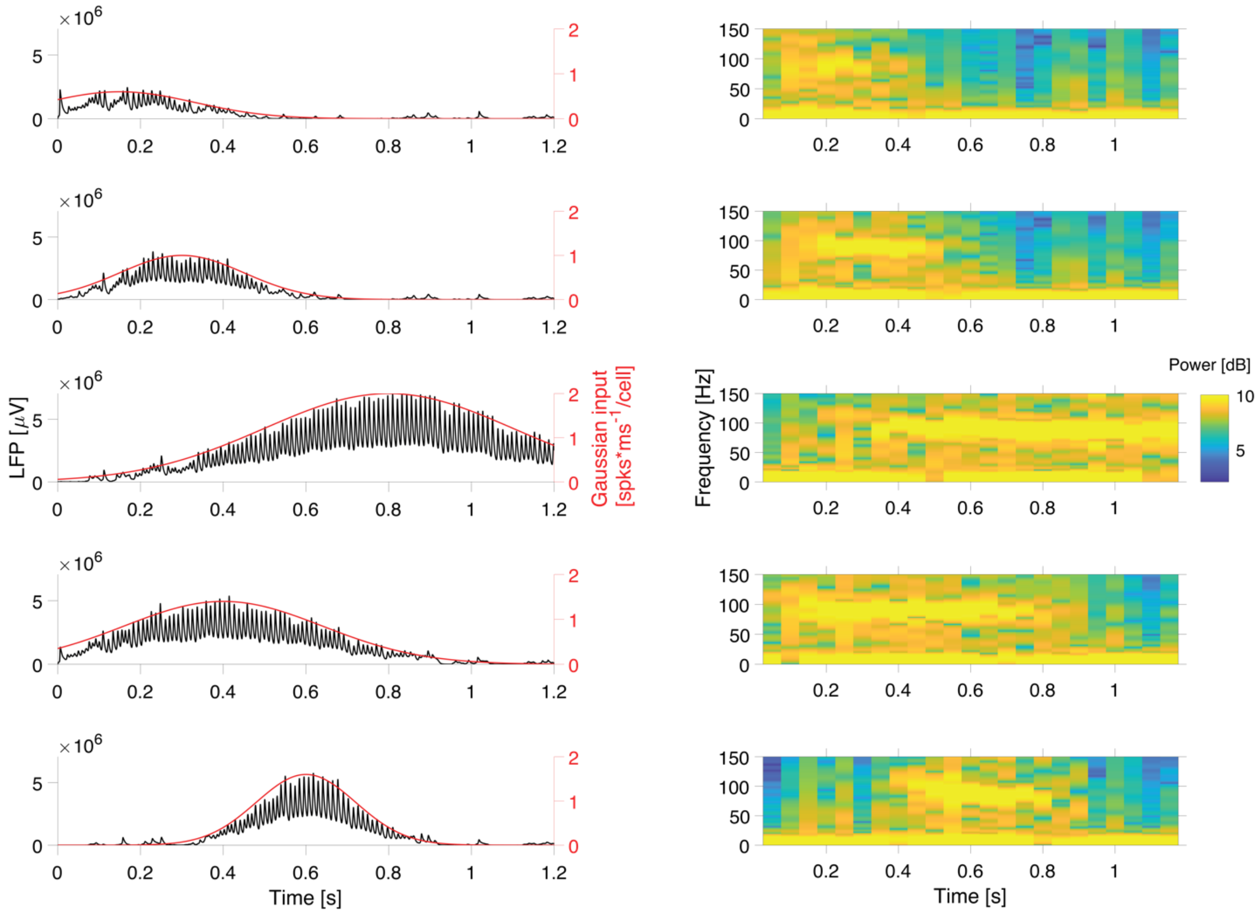


**Supplementary Fig. 9 | Gaussian input in the source generates different patterns of event-locked transient activity.** Exemplary time-frequency representation of the neural source fed by a gaussian input (red line). The duration and the temporal focality of the broadband $\gamma$-power activity can be tweaked by manipulating the mean (time of peak modulation) and the fullwidth at half maximum (modulation duration) of the gaussian function.

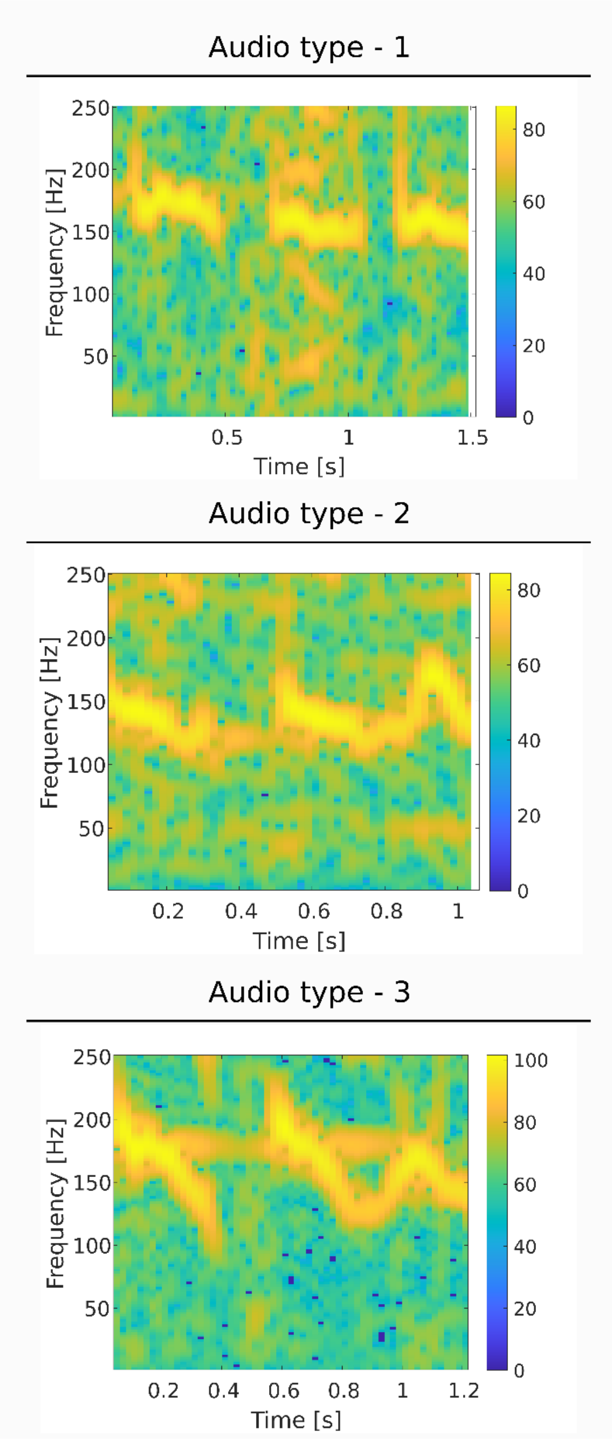


**Supplementary Fig. 10 | Spectrogram of the recorded audio used for the realistic scenario**. Spectrograms of different pitch patterns of utterances from three different participants.

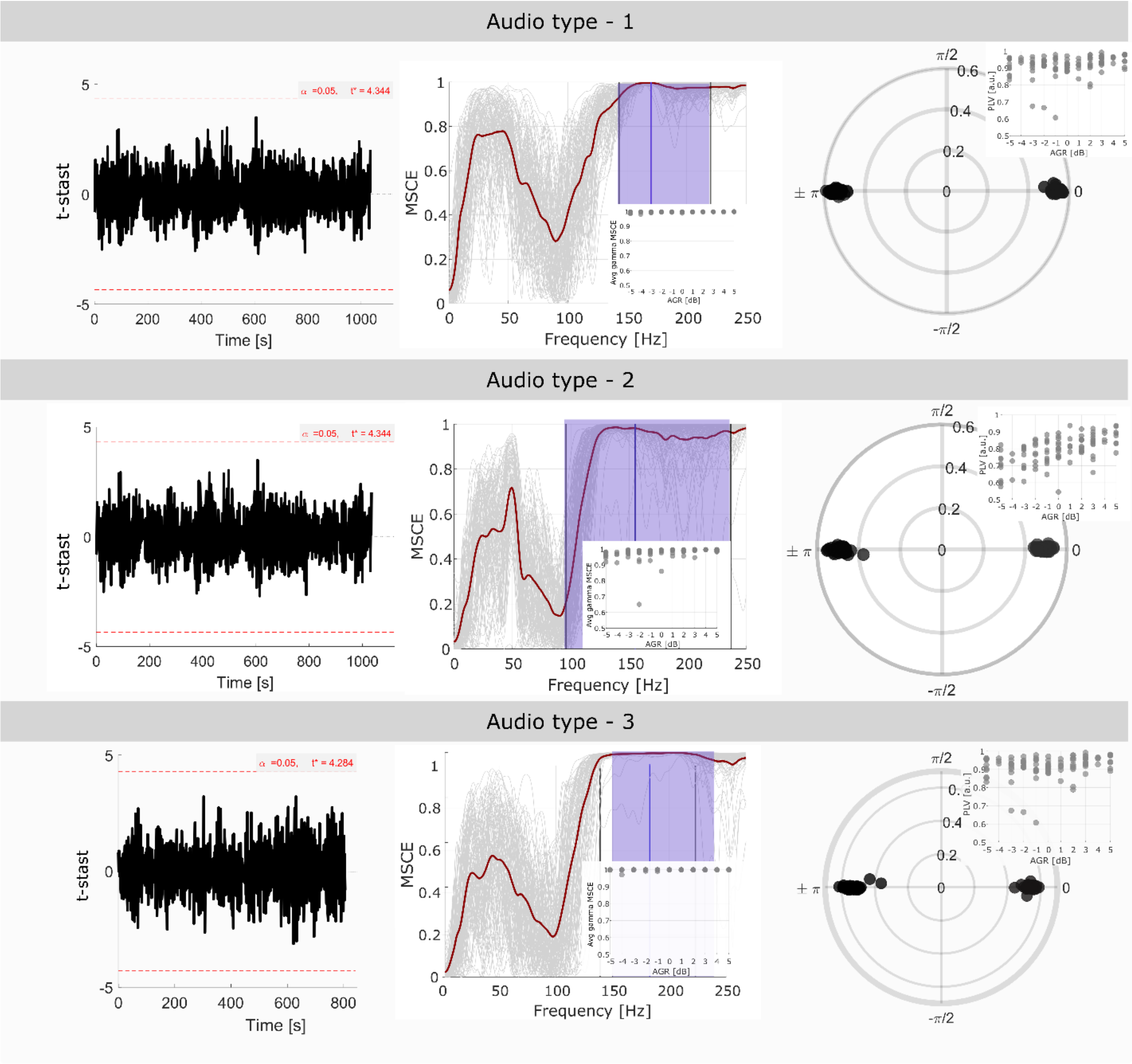


**Supplementary Fig. 11 | PCD perfectly retrieves the audio source in realistic simulated scenarios.** For every type of audio used during the recorded audio scenario (RAS) the PCD artifact estimation capacity is evaluated across trials in the time, frequency and phase domains. First column: one-dimensional statistical parametric mapping (SPM) used to evaluate statistical similarities at each sample point between the true and estimated source for each trial. No significant differences (t-stast values < t*) were found at any time point, indicating that across trials the true and estimated artifact source are identical from a statistical viewpoint. Second column: The magnitude-squared coherence estimate (MSCE) was computed between the true and the estimated artifact source. Gray lines represent individual trials. Mean MSCE across trials is denoted by the dark red line. The mean F0 across trials is shown as the blue vertical line, while the violet band indicates the possible range of speech artifact frequency band (SAFB) variations. The mean MSCE value in the SAFB was always above 0.97, regardless of the AGR of the trials. Third column: Phase differences between the true and the estimated artifact source. Each dot represents a trial phase difference. Estimated sources were either in phase or anti-phase relationship to the true source. Figure on the right corner aggregates the phase-locking value (PVL) across simulated trials at different AGR.

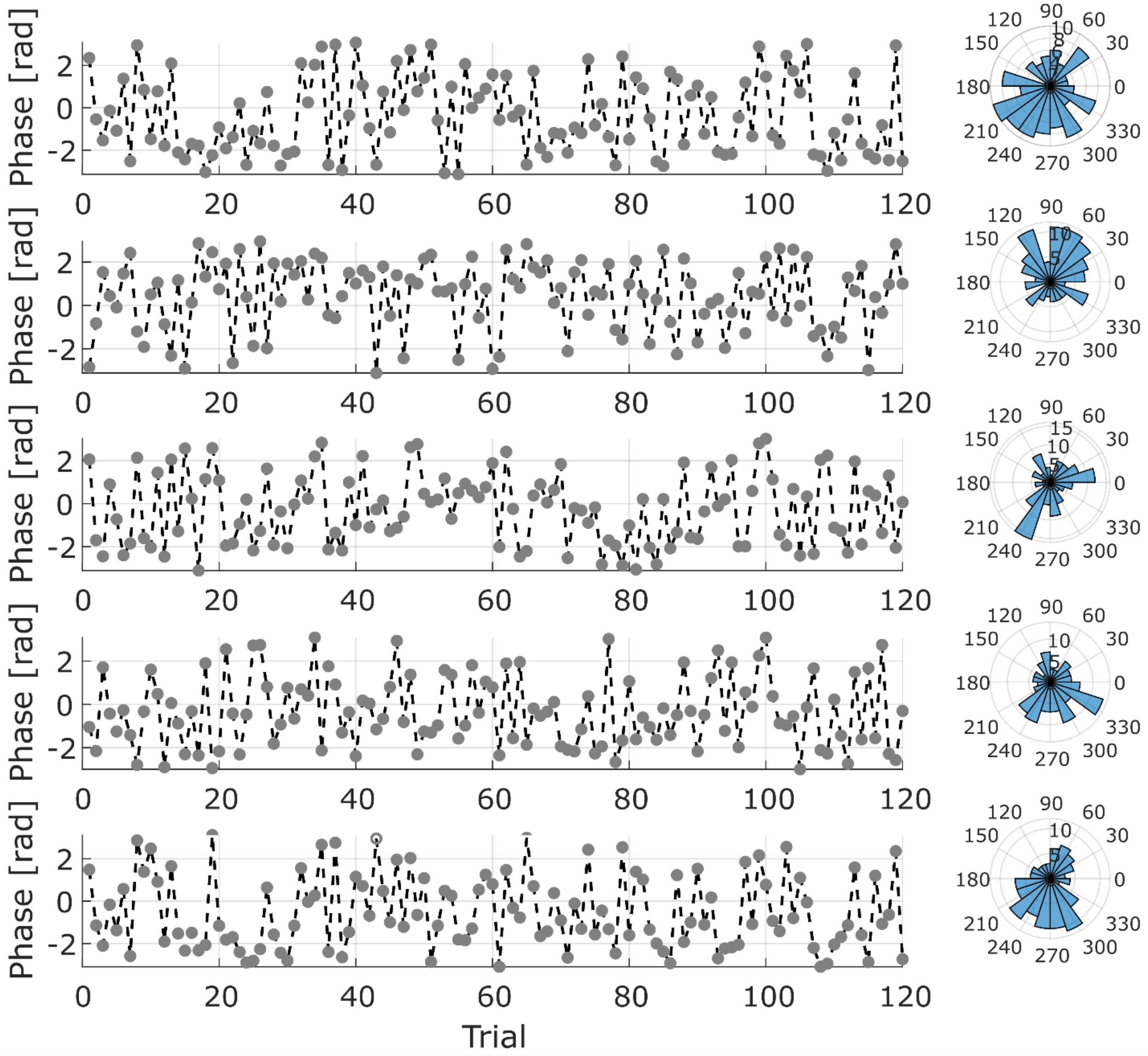


**Supplementary Fig. 12 | Artifact phase relationship varies across trials.** For a given participant and 5 random selected channels, the mean phase relationship between the recorded audio and each channel is shown for each trial. Figures on the right aggregate the distribution of the mean phase values across trials.

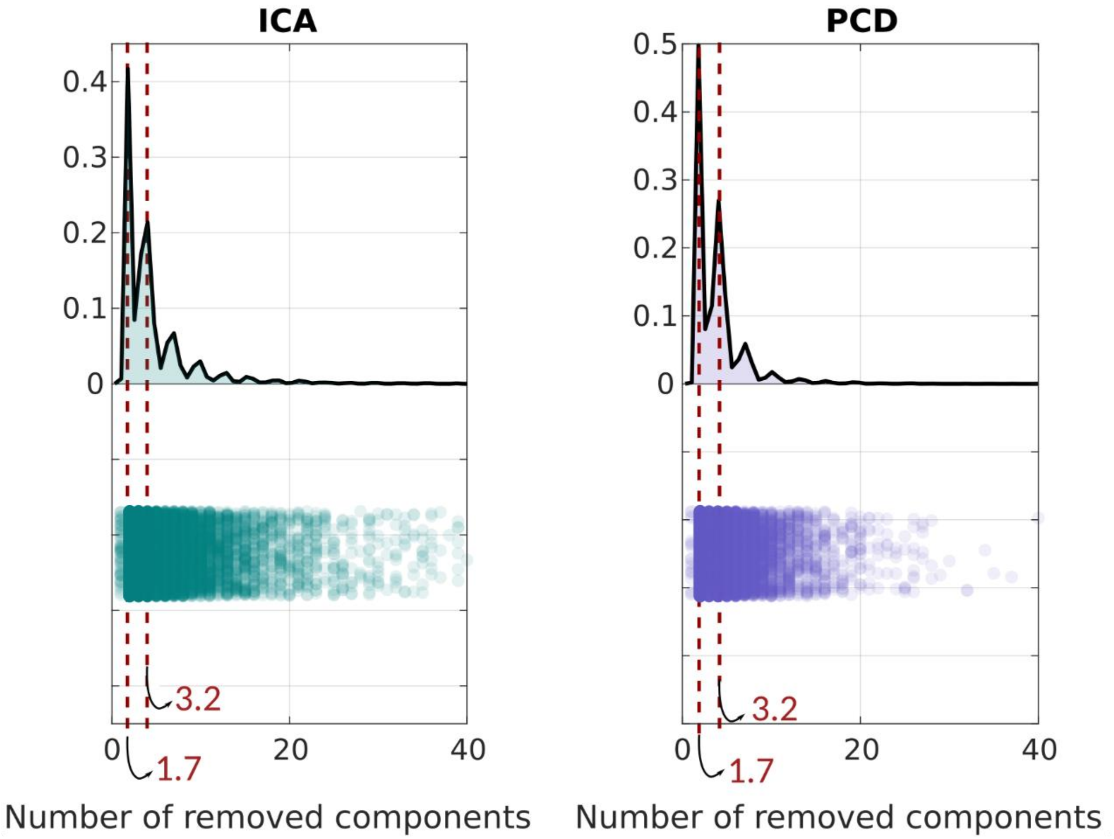


**Supplementary Fig. 13 | Number of removed components by ICA and PCD.** For each subject at each recording session, the number of removed component as automatically selected by ICA and PCD is drawn. Both methods in most of the cases removed around 1 to 4 components.
